# The trade-off between fractal morphology and function of Drosophila neurons

**DOI:** 10.64898/2026.10.14.759084

**Authors:** Ildikó-Beáta Márton, Balázs Péntek, Mária Ercsey-Ravasz

## Abstract

Studying correlation between structure and function in brain networks has long been a central and compelling topic in neuroscience. Using the FlyWire project’s fully digitized *Drosophila melanogaster* brain map, we analyzed a total number of 125330 3D skeletons of neurons from various functional super-classes and classes. By separating axonal and dendritic arbors of individual neurons we compare several morphological and topological properties, including cable length, fractal dimension, number of nodes, maximum width, and tree height. The fractal dimensions of neurons can reach up to 2.4, with dendritic arbors generally more complex than axonal ones, except in the afferent flow, where the situation is reversed. Correlating fractal dimension with cable length reveals that neurons from the same classes cluster in state space. These clusters densify and separate more clearly as we go deeper down the hierarchical annotation schema, indicating that the brain specializes fractal morphology for distinct functional classes.

## Introduction

The brain is organized across multiple structural and functional scales. Evidence suggests that fractal organization is present throughout this hierarchy, ranging from the microscopic architecture of neurons to the macroscopic arrangement of functional regions ^1^. In parallel, scale-free dynamics has been reported across a wide range of neural signals and spatiotemporal scales; however, a precise mechanistic account of how fractal brain morphology shapes neural dynamics remains to be established.

On the macroscale the most prominent geometric property is gyrification, the fractal-like folding of the cerebral cortex ^1^. It is still an open question if this has any functional role, it has been shown that the physical constraints during brain development alone suffice to produce such kinds of structures ^2^. The most studies related to fractal morphology were performed on the level of neurons because of the obvious fractal shape of their dendritic and axonal arbors ^3,4^. Especially the functional role of dendritic trees has been debated. Tosevski et al. measured the fractal dimension of 14 2D images of Golgi-stained pyramidal neurons from human basolateral amygdalae ^4^ finding significant differences between the fractal dimensionality of dendritic arborizations of two cell subtypes. More recently 3D images have also been studied, for example in ^3^ dorsal CA1 pyramidal cells using confocal microscopy, again fractal dimensionality was used to characterize dendritic arbors, and their conclusion was that neurons with higher dimensionality are more costly so the brain must make a trade-off between connectivity and energy expenditure. There have been several other studies ^5–9^, a more detailed history can be found in ^1^. However, most studies concentrated on measuring fractal dimensions of dendritic arbors in mammals, usually using a small cohort, and comparing a small number of cell-types. There was also an interesting study that managed to compare the fractal dimensions of dendritic and axonal arbors in Purkinje cells in rats drawing the conclusion that dendrites are more complex^10^.

Recently the whole brain of the female adult Drosophila melanogaster (fruit fly) has been mapped ^11–15^, the database containing a great amount of information about the approx. 140,000 neurons and 50 million synapses. Neurons in the fruit fly are of different shape than in mammals: the soma is in a layer on the periphery of the brain, disjoint both from the dendritic and axonal arbor ^12^. Starting from the soma, a primary neurite penetrates into the interior of the brain, where it then branches into two parts: dendrite and axon. Nevertheless, this large database provides an excellent opportunity to perform a wide-range computational study measuring the fractal dimensions and other morphological and topological measures of all neurons and their axonal/dendritic arbors and make comparisons between them. We also have information about the function of neurons: flows, super-classes and classes. To our knowledge this has not been performed yet for this database.

The pipeline of our study is the following: 1) we extract the 3D skeleton of neurons from the database; 2) a splitting is performed that separates the neuron in 4 parts: the soma with the primary neurite which is not interesting for our studies; the dendritic arbor, the axonal arbor, and the main axon (linker) connecting these two arbors (Fig. 1A). 3) We measure the total cable length of the separate parts: main axon, dendritic arbor, axonal arbor, and full neuron. 4) We also measure the fractal dimensions of full neurons, axonal and dendritic arbors. 5) From the skeletons we build the tree graphs and measure several topological properties of these, such as number of nodes, maximum width, and tree height. 6) All along these steps we will plot the distributions of properties for different functional super-classes and classes, we will compare axons with dendrites.

**Figure 1:**
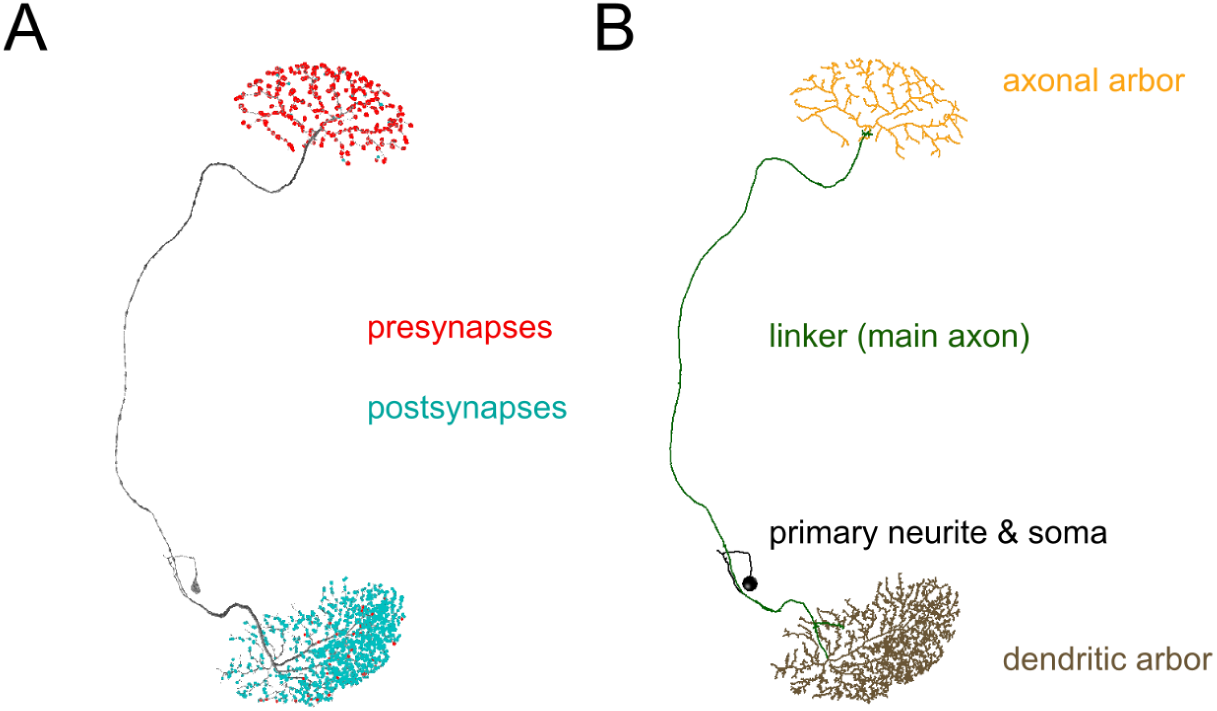
Example of Drosophila neuron. A) Neuron shown with its synapses. In general for the Drosophila, synapse types are more mixed on the neuron arbors ^12,29^. B) Skeletonized neuron split based on synapse flow centrality. Generally, Drosophila neurons consist of 4 compartments: primary neurite (black), linker/main axon (green), dendritic arbor (brown) and axonal arbor (yellow).

We will show that main axons (linker) have an exponential cable length distribution, being in agreement with our previous measurements ^16^, and also similar to findings in mammals ^17,18^. However when focusing on fractal-like structures of dendritic and axonal arbors lognormal distributions appear. This is expected because of the multiplicative branching process in the arbors ^19^. The total cable length distribution of neurons is given by the combination of these two.

We will compare properties of axonal and dendritic arbors, and we will show that the fractal dimension, at least here in the Drosophila, is not always larger in dendrites. The exceptions are the afferent super-classes (incoming neurons), where the axons show higher complexity on average.

In general, we will examine how fractal dimensions and various morphological/ topological properties of neurons differ across functional classes. As we delve deeper into the functional hierarchical classes of neurons, these properties - and the correlations among them - become increasingly effective at distinguishing the classes. In other words, the brain appears to specialize in producing fractal morphologies for specific functional types.

## Results

### The Drosophila Database

The fruit fly (Drosophila melanogaster) is a useful model organism in biology from a neuroscience standpoint, due to its rich functionality. In addition to basic functions like vision, flight, and walking, the fruit fly exhibits more complex behaviors such as courtship and aggression ^20–26^.

Recently, a milestone in connectomics was reached with the FlyWire project, which mapped the whole-brain connectome of Drosophila using automated image analysis on previous electron microscopy recordings ^15^. A large number of scientists from all around the world collaborated in the proofreading and annotating of 139 thousand reconstructed neurons. Therefore, each neuron’s precise location and morphology became available. Some of the functional properties are also provided in the form of functional roles/classes organized in a hierarchical structure. The neuron’s ‘flow’ indicates the direction of information/signal flow: intrinsic neurons operate completely within the brain, while non-intrinsic neurons can be of afferent and efferent types projecting inward and outward. Each ‘flow’ is subdivided into ‘super-classes’: intrinsic → central, optic, visual projection and visual centrifugal; afferent→sensory, ascending and optic; and efferent → descending, motor and endocrine respectively. In the case of these non-intrinsic neurons there are many cases when dendritic or axonal arbors are not available, that is the main reason why we analyze a lower number of neurons, 125330. Some of the ‘classes’, such as the sensory ones are also mentioned in later sections of this study, and a more comprehensive discussion of this hierarchical annotation can be found in the literature ^14^.

In addition to neurons, the dataset contains approximately 50 million synapses predicted with high accuracy ^11,13,27^, together with the predicted neurotransmitter types ^28^. The project’s open access web and python interfaces (Codex and fafbseg.py) made it possible to analyze the fractal properties of neurons and their arbors, and their relationship to functional roles at an unprecedented scale.

Our aim is to analyze the full neurons and its dendritic and axonal arbors separately as well. For this, we need a method to split the neurons into compartments based on the synapse locations. We chose the algorithm based on synapse flow centrality developed in a previous study ^29^. Generally for the Drosophila, the splitting results in 4 parts: primary neurite, linker (main axon), dendritic and axonal arbors (illustrative example shown in Fig. 1. However, it is worth noting that there are some cases where the splitting algorithm yields inaccurate results, partly because of the mixed nature of pre- and postsynapses on the Drosophila neurons. Some examples for atypical neurons and splitting results are discussed in more details in the Methods section.

After extracting the skeleton of neurons and splitting them in 4 parts, we measure the total cable length (*L*) of the full neurons, main axons, and the axonal/dendritic arbors (see in Methods section). Based on the skeletons we also want to measure the fractal dimensions of full neurons, axonal and dendritic arbors. Because of the huge number of neurons we needed to make a compromise between applying computationally efficient and highly precise methods. We applied the box-counting method ^30^. Having to measure the fractal dimension (*D*) for all 125330 neurons we had to automatize the algorithm (See in Methods section). Measuring for the whole neurons the algorithm works well, but the splitting sometimes identifies very small non-fractal-like structures for dendrites or axons (Supplementary Figure S1.5). The dimensionality of those with the automatized algorithm gives some numerical errors (*D*<1). Actually there are more reasons why errors may appear in the fitting making the fractal dimension go below one: 1) we are working on the skeletons of the neurons not on the true images, this always decreases the dimensions, especially when they are close to one; 2) the distribution of points is not homogenous in the skeleton and this can affect the measurements of *D* especially in case of small parts; 3) There are few exceptional cases in this database where some errors happen during the skeletonization of the reconstructed neurons, resulting in missing nodes. To avoid fragmented neurons in the database, a healing process is applied and the missing parts are replaced by unrealistically long straight lines (example shown in Supplementary Figure S1.2). In order to omit these neurons from our analysis, we employ a thresholding method based on the distances between neighboring nodes in the skeleton, a more detailed description can be found in Methods section; 4) That small percentage of axonal and dendritic arbors for which we obtain *D*<1 usually has very small cable length and maximum width (we will show below in the results), meaning these are highly elongated structures where the branching density is low. We introduced some methods trying to correct these errors (see Methods section) still a certain percentage of axons and dendrites remained with *D*<1 (2% intrinsic axons, 1.77% intrinsic dendrites, 8.57% non-intrinsic axons, 16.51% non-intrinsic dendrites). However, our main goal here is not necessarily the precise measurement of the fractal dimensionality, but rather the comparison of different functional classes of neurons, and as mentioned above, these errors come mainly from the splitting procedure and obtaining extremely small non-fractal like structures that are analyzed after that during the automatized algorithm.

Besides these measures we also wanted to measure some topological properties of the full neurons and dendritic/axonal trees. The skeleton itself is not a simple tree, there we have many more nodes included in the graph trying to follow the curvature of the cables of the neuron, as seen in Fig. 1, and explained in Methods section. Neuronal skeletons were downsampled ensuring that all ‘middle nodes’ were removed, to reduce node density along branches while preserving branching structure and connectivity. From the original skeleton we build up the tree of the full neuron, the root of which will be the soma. The tree of the axon and the dendrite are the two subgraphs with roots starting at the nodes identified by the splitting algorithm. These trees are not always binary trees, quite often axons produce 3-ary branches ^31^. We focus on the number of vertices (nodes) in the tree (*N*), the number of levels, the height (*H)* of the tree, and the maximum width (*W*) of the tree, which is the maximum number of nodes at any level (Methods section, Figure S1.6).

### The distribution of cable lengths of different parts of neurons

There have been several previous studies related to discussing the distribution of cable lengths and especially the axon length of neurons, so first we start with clarifying this issue. It has been observed first in the white matter of the macaque ^17^, later in mice ^18^ that the distribution of the axon length crossing the white matter between different brain areas shows an exponential distribution, called the ‘exponential distance rule’ (EDR). Based on this the EDR random network model has been applied as a null model in all these mammals and has been shown to explain a lot of the topological properties of the inter-areal cortical network of those species. In a recent study we checked this also in the Drosophila ^16^, by measuring the distribution of the main cable between the axonal and dendritic arbors, we proved the validity of the EDR and also the EDR model on the level of neuropils (functional regions). In that study the starting point of axonal/dendritic arbors were identified slightly differently, here we use the splitting algorithm of ^29^. Nevertheless the result is very similar, the distribution of the cable length of the main axons shows an exponential decrease (Fig. 2A). Looking at the dendritic and axonal arbors, however, we see more like lognormal distributions of the cable lengths (Fig. 2B). This is explained by the multiplicative branching process in these fractal-like tree structures ^19^. However, the total cable length of the full neurons (Fig. 2C) must be somehow a combination of the long axon cable, and the axonal/dendritic arbors, resulting in a more distorted distribution (Fig. 2C) deviating from a simple lognormal. This interesting difference between the distribution of main axons and local fractal arbors shows how the brain optimizes in a different way global vs local communication. Main axons have relatively few synapses in the Drosophila, their main role is to transmit information over long distances, even to the other side of the brain if necessary. This is achieved through the EDR rule, as observed also in other species. At local level high connectivity is needed, which can be achieved most efficiently by growing these fractal-like tree structures, the dendritic/axonal arbors that develop synapses and form connections between them.

**Figure 2:**
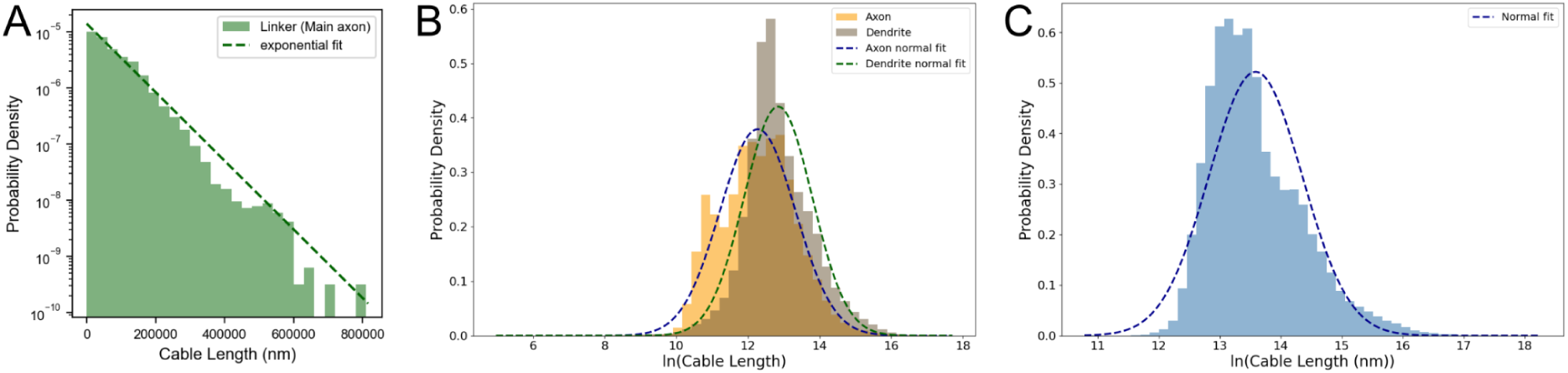
Distribution of cable lengths values. A) For main axons; B) dendritic (grey) and axonal (yellow) arbors and C) full neurons. Dashed lines indicate fittings with A) exponential B,C) normal distributions.

### Comparing properties of axonal and dendritic arbors

Most of previous studies concentrated on studying dendritic arbors, these being more easily available in mammals. Here we were curious about comparing axons and dendrites in different functional classes of neurons. In Fig. 3A we plot the distribution of fractal dimensions of full neurons, axonal arbors and dendritic arbors. At this first step we analysed only intrinsic neurons, having 114190 of them. As we mentioned when describing our numerical method, when applying it on the full neurons it works very well, only for 9 neurons the fractal dimension goes slightly below 1. In the case of axons and dendrites however, because of the issues described in the Methods section there are many more errors. Usually these are extremely small parts of the neuron, that are identified by the splitting algorithm as the axonal/dendritic arbor, there is no fractal structure, the cable length and branching is small, so our box-counting algorithm fails and the fitting provides a dimension smaller than 1 (see Supplementary Figure S1.4 for an example). However, these are not representative of the fractal properties of neurons, and represent a small part of the sample: 2% of axons and 1.77% of dendrites. We ignore these, they will appear on the plots, we will most of the time draw a red line at *D*=1 at the distributions, or on the violin plots. We can see that there is a significant difference (p<10^-10^) between the distribution of dimensionalities for axons and dendrites, dendrites having larger dimensionalities. Looking at this total cohort of intrinsic neurons all other properties - the number of nodes in the trees (Fig. 3B), the height of trees (Fig. 3C) and maximum width of trees (Fig. 3D) - show a similar behavior, namely dendrites are larger and wider than axons.

**Figure 3:**
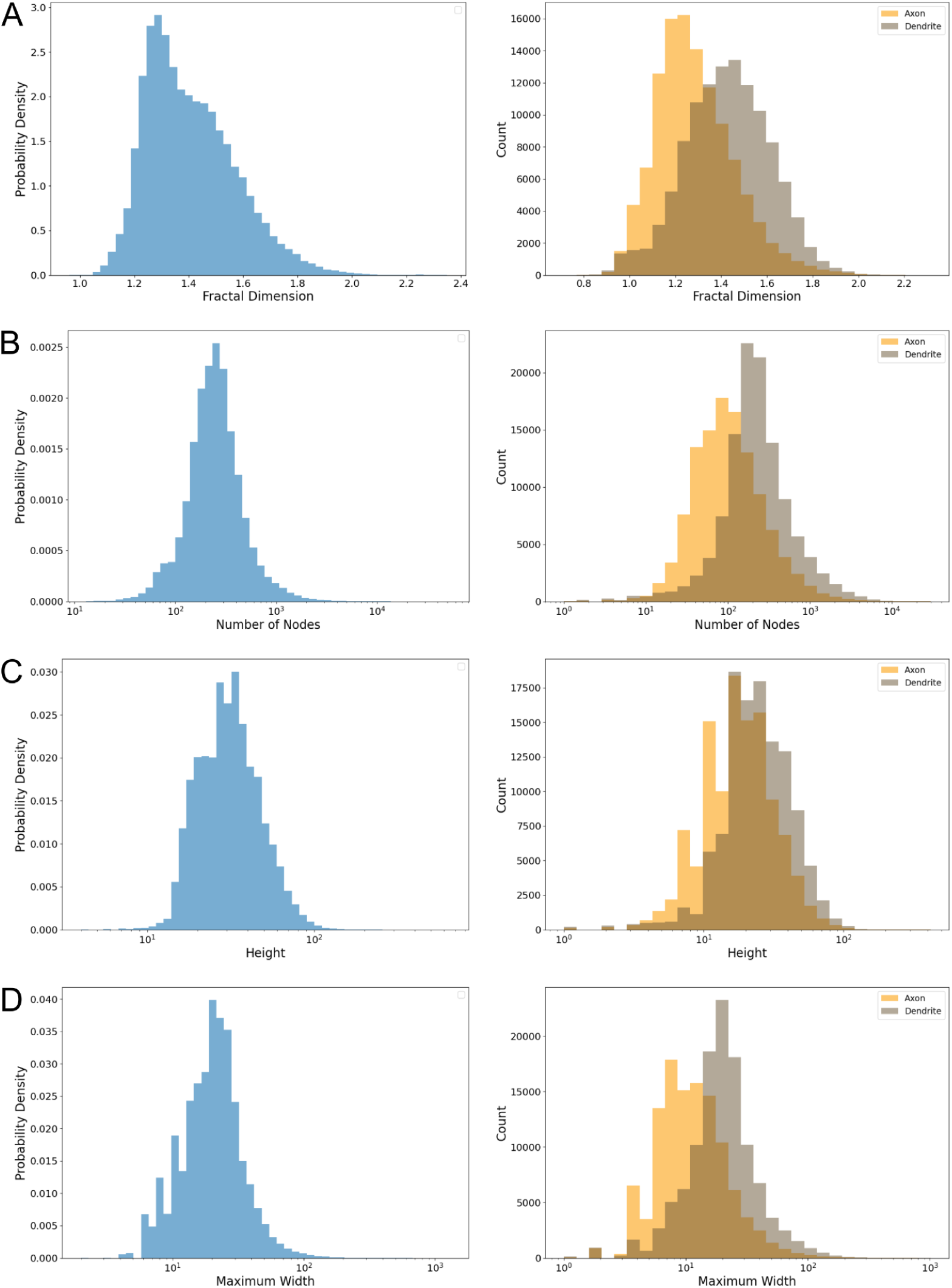
Distribution of different properties for the full intrinsic neurons, and their comparison between axons and dendrites. A) Fractal Dimension, B) Number of nodes, C) Height of the tree, D) Maximum width (p-values are <10^-10^ for all 4 comparisons).

Having information about their functional role, neurons being classified in super-classes and classes, we wanted to study how these properties may differ between these classes. High fractal dimensionality requires extra energy for the brain, some previous studies suggested already that a trade-off between high dimensionality and energy expenditure must be made by the brain when developing different cell types ^3^. In Fig. 4A we show with violin plots a comparison between the distribution of fractal dimension of axonal/dendritic arbors of different intrinsic super-classes: optic, central, visual projection, and visual centrifugal. Above the violin plots there are the distributions for the full neurons. Fig. 4B shows the distributions of cable lengths, Fig. 4C the maximum width of the trees in a similar arrangement. In Supplementary Figure S2.1, S2.2 we also show the number of nodes and the height of the trees. For the comparison of distributions of axons and dendrites we used Cliff’s δ as the effect size metric^32^ (see Methods section). The values appear under the violin plots. These properties are also expected to correlate with each other, as seen in Fig. 5.

**Figure 4:**
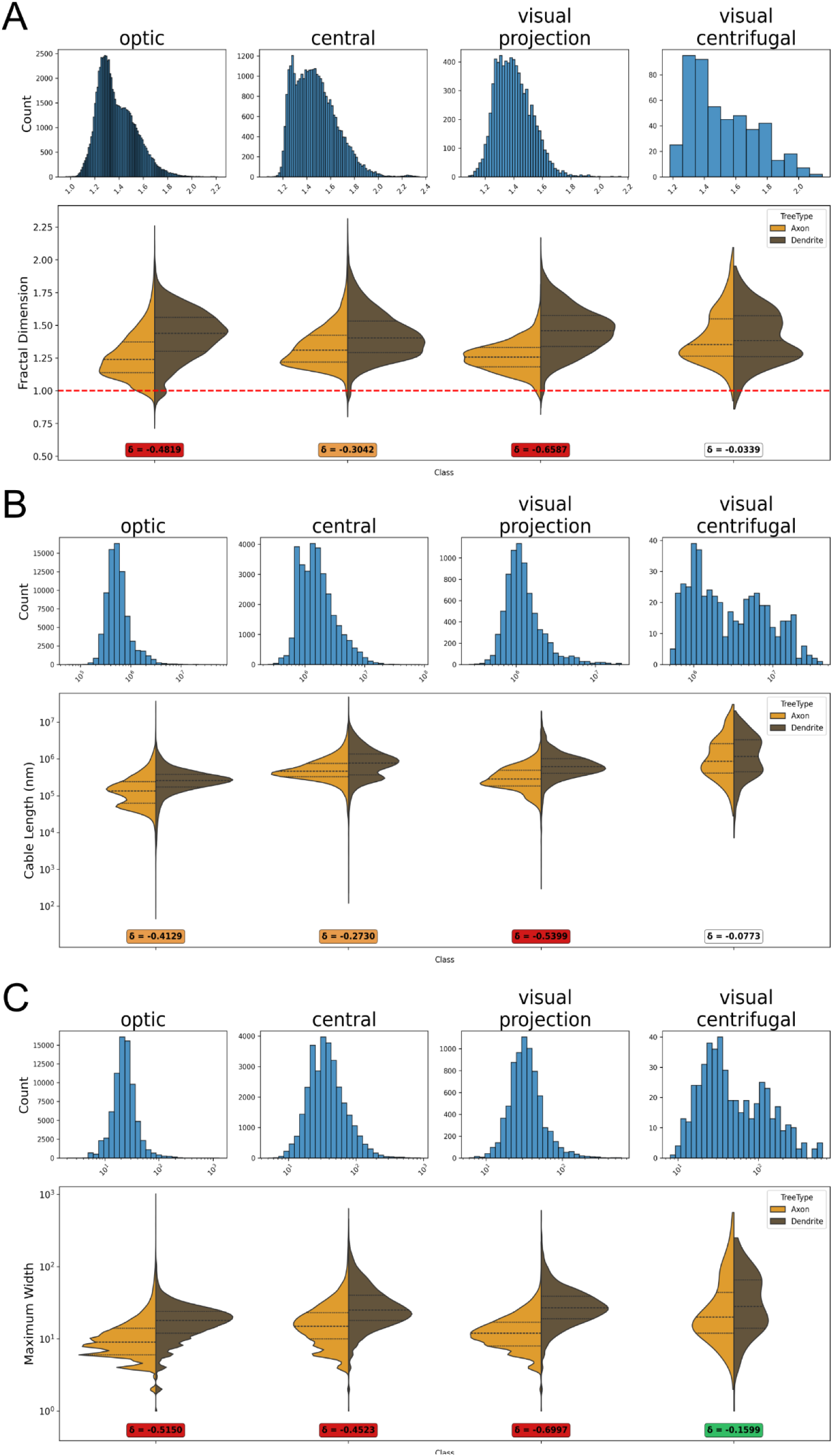
Comparing the 4 super-classes of intrinsic neurons. We show the histograms of the properties for the full neurons and violin plots comparing the distributions for axons and dendrites for the A) fractal dimensions, B) cable lengths, C) and maximum widths. The Cliff’s δ effect size values for the comparison of all neuron properties are shown in Supplementary Table S2.1.

**Figure 5:**
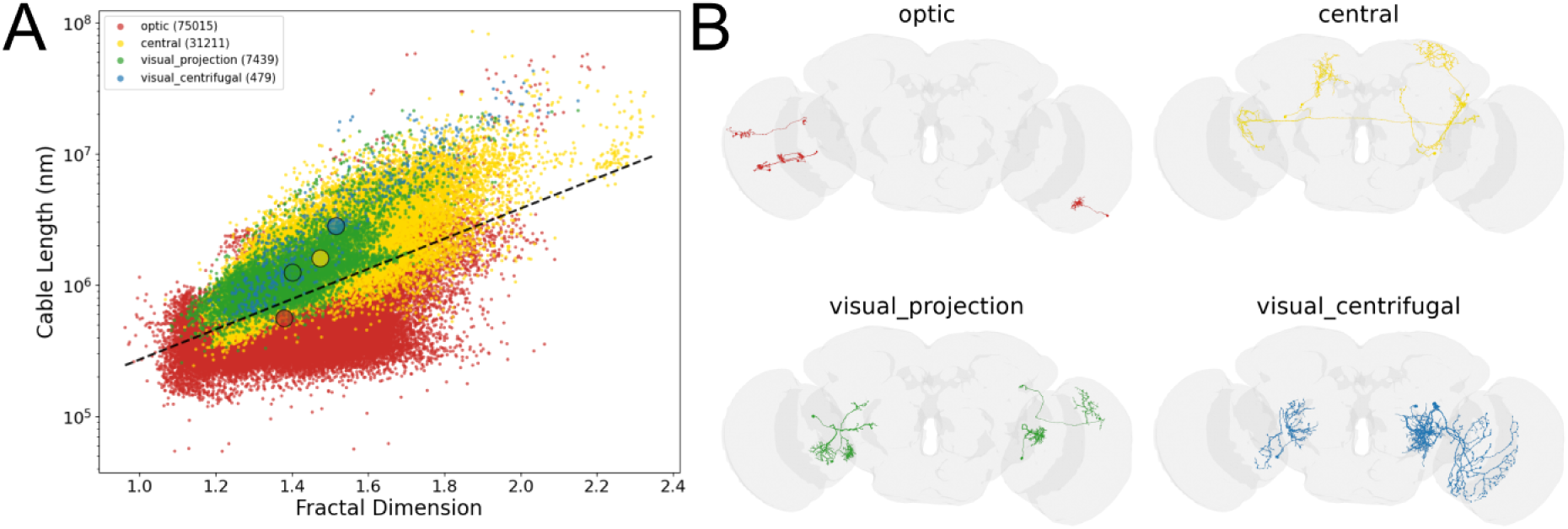
Intrinsic neurons. A) Colored correlation plots between fractal dimension and cable length, B) 4 typical neurons from each intrinsic super-class shown in the fruit fly brain. See the correlation plots between fractal dimension and number of maximum width/nodes/height in Supplementary Figures S1.3 - S1.5. The large colored circles represent the centers of mass.

For example in the largest, optic super-class (red in Fig. 5) with 75015 neurons the fractal dimension can vary over a wide range from 1 to 2.2. While the majority of neurons have a smaller fractal dimension (below 1.6) and smaller cable length and maximum width, there is a tail of the histogram going up to these large values. The best visualisation of these super-classes and their properties is in Fig. 5, where we colored each dot based on the function of the neuron and we show the correlation between the fractal dimension vs cable length. We can see how the red cloud is the largest one starting from the smallest properties, but still some red dots can be seen at very high cable length and large fractal dimension values (as also indicated by the histograms). Then the central super-class (yellow) follows with 32211 neurons, the cloud being shifted a bit upwards, this is the super-class that reaches the highest fractal dimensions from all, going up to 2.4. Even an interesting separated little cloud can be noticed at high values. The visual projection (dark green) super-class contains 7439 neurons, the cloud overlaps the central yellow cloud, however the correlation between properties seem to get much stronger, the cloud is a bit thinner. The fractal dimensions are not so large in this super-class, only a few neurons go above 1.75. The visual centrifugal (blue) super-class contains only 479 neurons. We can see this cloud to be even thinner (more correlated) and slightly upwards and to the right in Fig 5A, meaning these neurons have larger cable lengths and larger fractal dimensions as also shown by the histograms. The fractal dimension again spans a broad range, some neurons take values above 2 (number of nodes, tree height shown in Supplementary Figure S2.1, S2.2). These being functional super-classes they still contain a lot of different types of neurons, so it is somehow understandable or even expected that these large clouds overlap. In order to distinguish them more easily we show their center of mass with larger colored dots (see Methods section). The black line in Fig. 5A is a linear fit over all points taking the logarithm of cable length values and the fractal dimension. Deviations from this line could tell us about the scale of global and local communication of a given neuron. One with a lower fractal dimension and high cable length could indicate a neuron performing a more global information transfer in the brain, with a long main axon (linker) and simpler dendritic/axonal arbors. On the other hand, high fractal dimension and small cable length could indicate a stronger role in local communication with complex dendritic/axonal arbors. The complexity of different super-classes is illustrated in Fig. 5B by plotting 4 typical (closest to the center of the cluster of the neuron super-class) neurons for each class on the brain map.

Comparing the distribution of properties of axons and dendrites we can notice that in the optic, central and visual projection super-classes the dendrites are much larger, showing high fractal dimension, cable length etc. (Figure 4). In the visual centrifugal super-class this is still true, however, the difference is much smaller. As shown in the next chapter we also studied the non-intrinsic super-classes where we were able to analyze the afferent/efferent flows (incoming/outgoing neurons). This question is interesting because in previous studies in mammals mainly dendrites were studied, very few results are available about comparing fractal properties of axons and dendrites, the most important study concludes that dendrites are more complex^10^. Nevertheless, we must remember that also in mammals there are lots of types of neurons. Until now our findings show that all 4 super-classes of intrinsic neurons in the Drosophila support the same conclusion.

### Afferent and efferent neurons

In the case of non-intrinsic neurons we have the following super-classes: sensory, ascending, optic – in the afferent flow (incoming neurons), and descending, endocrine, and motor in the efferent flow (outgoing neurons). Here we can make several interesting observations. In all 3 afferent super-classes axonal arbors are more complex on average (Fig. 6A), while in efferent super-classes dendrites are more complex on average (Fig. 6A). It seems like arbors inside the brain have more complex structure than those outside, which is maybe because outside the brain the constraints of development are stronger. The second interesting observation is that the two flows are almost perfectly separable on the correlation map in Fig. 6B, indicating that the afferent neurons are less complex, with shorter cable length and smaller average fractal dimension. In Fig. 6C we show 4 typical neurons from each super-class, with the increase in complexity being visually apparent.

**Figure 6:**
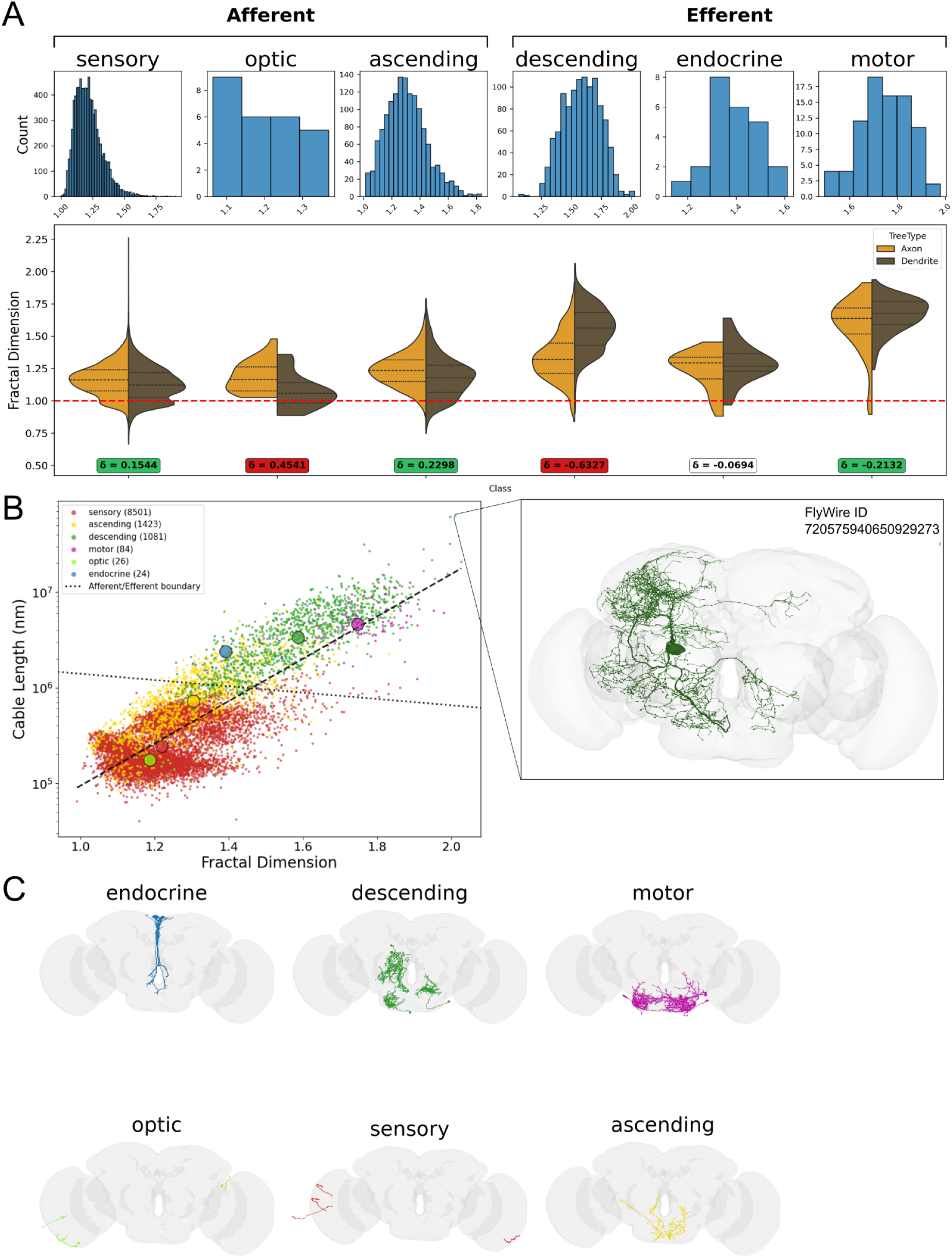
Extrinsic neurons. A) Histogram of fractal dimensions of extrinsic neurons. Violin plots for the distributions of the fractal dimension of axonal and dendritic arbors. The Cliff’s δ effect size values for all neuron properties are shown in Supplementary Table S2.2. B) Correlation between the fractal dimension vs cable length in extrinsic neurons, dots being colored based on the super-class of neurons (see legend). See the violin plots and correlation maps for other properties in Supplementary Figures S2.6 - S2.11. The large colored circles represent the centers of mass. On the right we show the 3D skeleton of the descending neuron with the longest cable length (∼6.5 cm) positioned inside the Drosophila brain. C) 4 typical neurons from each afferent/efferent super-class shown in the fruit fly brain.

Besides being able to separate the two flows: afferent and efferent, looking at Fig. 6B we notice again how neurons from the same super-classes are generally grouped together, meaning they have similar morphological properties. In this state space of morphological properties these functional super-classes start to get separated. The few neurons in the optic super-class (light green) have the smallest fractal dimension and length, after that follows the sensory super-class (red) in which most neurons have cable below 10^6^ nm and fractal dimension below 1.5, still we find a few neurons with dimensions even above 1.8. The ascending super-class (yellow) is slightly shifted above the red to longer cable length values, but similar fractal dimensions. The most surprising separation occurs by the descending supper-class (dark green), that has almost no overlap with the sensory super-class, being shifted to much larger cable length and fractal dimension values. This is where the afferent and efferent flows are separated. The total cable length of some of these neurons can reach ∼6.5 cm (see the 3D skeleton of the neuron in the right side of Fig. 6B), because of their complicated fractal structure, which is surprisingly long compared to the size of the fruit fly. The endocrine super-class (blue) contains very few neurons lying along the upper edge of the green cluster, and the other interesting super-class is the motor (violet) that again separates itself in the upper corner, at not so large cable length values, but rather large fractal dimensions.

These are functional super-classes that still contain many more specialized neuronal classes and subclasses, but we can already notice how a trade-off between the costly morphological properties and the function of neurons have been achieved during development.

### Afferent sensory neurons

If we go even deeper in the hierarchical set of annotations studying the non-intrinsic sensory super-class for example, and the classes included in it: mechanosensory, gustatory, olfactory, visual, hygrosensory, thermosensory and unknown sensory, we can notice a similar behaviour of the distributions as until now (Fig. 7), however on the correlation map of the fractal dimension and cable length the colored clusters take up even smaller areas, showing that a functional class gets to be even more specified by these morphological properties (Supplementary Figures S2.12 - S2.17). The visual class (blue) forms a well-defined triangle-shaped cluster in the lower left corner with relatively small fractal dimension and cable length values. Above it we can see as a red patch the mechanosensory neurons grouped together, although a part of this class is a bit scattered at higher cable length values. The olfactory neurons (dark green) again form a very nice cluster at relatively high cable length values and average fractal dimension 1.3, as also shown by the histogram, but a separate, less denser cluster can be seen with a higher fractal dimension (indicated by the long tail of the histogram) but lower cable length. It seems like a different subclass of these olfactory neurons. The gustatory neurons (yellow) form a patch in the middle, but this cluster is a bit more elongated, going up to higher cable length values. The hygrosensory class (light green) is interesting because it shows two peaks in the histogram, and indeed we see the light green dots to be separated in two little clusters (seen to overlap with the left part of the dark green cluster and more scattered above the top of the yellow). There are only 29 thermosensory neurons (light blue), but we can see them grouped together relatively close in the upper corner. The unknown sensory (violet) class spreads around a larger area, but is definitely characterized by the largest fractal dimensions of all. Morphologically, on average, these unknown sensory neurons appear most similar to thermosensory neurons. However, determining their exact modality will require definitive experimental or molecular evidence. In Fig. 7B we show examples of 3D skeletons of neurons with the largest volume from each class.

**Figure 7:**
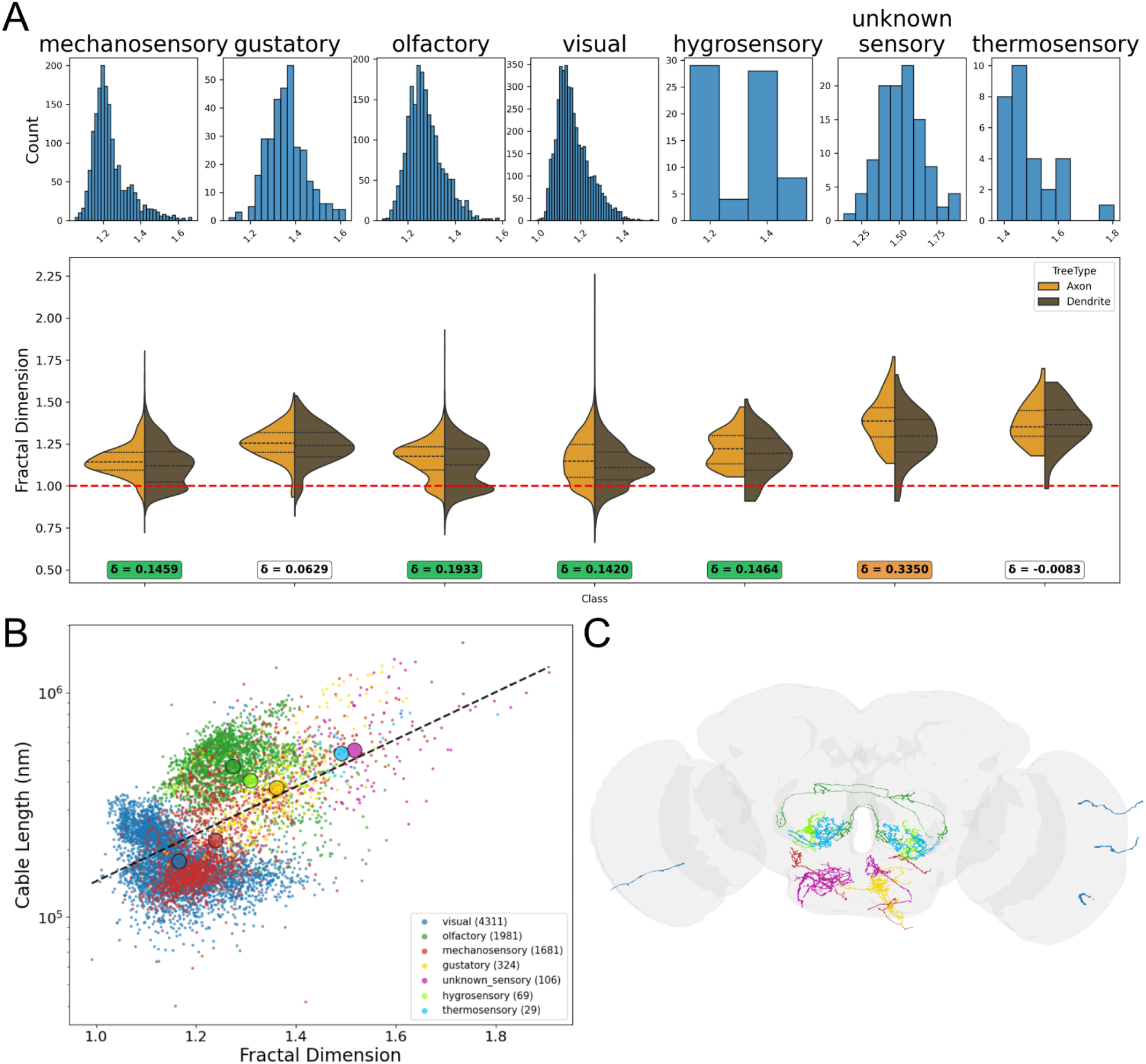
Afferent sensory neurons. A) Histogram of fractal dimensions of afferent sensory neurons. Violin plots for the distributions of the fractal dimension of axonal and dendritic arbors. The Cliff’s δ effect size values for all neuron properties are shown in Supplementary Table S2.3. B) Correlation between the fractal dimension vs cable length in afferent sensory neurons, dots being colored based on the class of sensory neurons (see legend). C) 4 typical neurons from each sensory class shown in the fruit fly brain. Neuron colors correspond to legend in B.

## Discussion

The trade-off between the necessary energy and efficient information transfer is a crucial question in the brain, not only on the level of the connectome but even on the level of single neurons. The different parts of a neuron for example, the main axon cable that usually has almost no (or much fewer) synapses vs the axonal/dendritic arbors that have the role of communication and synapse formation, have very different structures. The main axon is a more simple cable that sometimes needs to be able to project to extremely long distances (even opposite sides of the brain). Its length distribution follows the exponential distance rule ^16,17^. The axonal/dendritic arbors on the other hand need to transfer information locally with high efficiency by producing sometimes a huge number of synapses, which can be best achieved through arbor-like fractal structures on which the synapses appear in great numbers after connections are formed. The length distribution for these arbor-like structures is naturally different and can be approximated with lognormal distributions because of the multiplicative branching process characterizing these trees. Recently it has been hypothesized that cable length distributions of neurons follow lognormal distributions ^19^, and this indeed seems to be true for the axonal/dendritic arbors, but we must not forget that the total cable lengths of full neurons consists of the sum of the lengths of the main axon cable (exponential), the axonal and dendritic arbor and in the case of the fruit fly we would need to consider even the primary neurite.

It is also worth mentioning that most of the previous studies focused on measuring fractal dimensions of dendritic arbors in mammals ^3,4^. The reason is probably because in mammals dendrites are around the soma and are more easily accessible for study. There have been only a few studies comparing dendrites and axons in Purkinje cells ^10^, and the conclusion was that dendrites are more complex. Interestingly, looking at intrinsic neurons here we again see dendrites with higher fractal dimension and more complex topological properties, and this remains true even when we investigate them separately by super-classes. However, when looking at afferent and efferent flows the arbors within the brain (axons of afferent/incoming neurons and dendrites of efferent/outgoing) were found to have more complex structures. One possible explanation for this could be that constraints of development are much stronger outside the brain, so axonal/dendritic arbors outside the brain do not develop such complex structures.

Another interesting observation was that the afferent flow was overall much more simpler than the efferent one. These incoming (afferent) neurons just bring signals (mainly sensory inputs) into the brain, dropping off the signals at the low functional hierarchical level of the brain, which are then processed further on at higher levels. So these incoming ‘cables’ do not need to be so complex. The feedback process, however, needs to collect information from the brain in order to transmit outside what action needs to be performed (e.g. motor neurons, descending neurons etc.). This is a more complicated process that is performed by neurons with higher fractal dimension and longer cable length.

It has always been an important question if fractal morphology of dendrites or axons carries somehow an important functional role. We demonstrated here, especially on the colored correlation maps of fractal dimension vs cable length (Figures 6B & 7B), how neurons from the same super-class or the same class group together, meaning they have similar properties. When we plot super-classes these clusters are still very large, because we still have many different types of neurons in these large super-classes (Figure 5). As we get down the hierarchy many of these clusters seem to get smaller and denser, like they would be much more specialized by these properties. The clusters even start to separate from each other. It seems there are functions for which the brain needs complex neurons with high fractal dimensionality (e.g. efferent motor and descending), but there are functional classes for which simpler neurons also suffice (e.g. afferent neurons).

This study achieved for the first time a wide-range computational study of measuring the fractal dimension and other morphological/topological properties of 125330 Drosophila neurons extracted from the FlyWire database ^12^, comparing 4 different super-classes of intrinsic neurons, 6 different super-classes of afferent/efferent neurons and 7 classes of sensory neurons. Our main results show that as we delve deeper into the hierarchical classes of neurons, the morphological properties within a given class become more similar to one another, often forming clusters in state space. In essence, the brain appears to specialize in producing fractal morphologies tailored to specific functional types. Because neurons with higher fractal dimensions are metabolically more costly to develop, the brain must navigate a fundamental trade-off between energy expenditure and connectivity complexity. Ultimately, our results provide a quantitative framework that could help future studies more clearly delineate cell types and subtypes in mammalian databases, elucidating exactly why certain functional classes require high fractal dimensionality while others do not.

## Methods

All analyses were implemented in *Python* using the *fafbseg* (FlyWire) and *navis* libraries. Neuronal morphologies were analyzed by computing a set of topological (tree-based) and geometric properties.

### Preparing the data for geometric properties

First we loaded the skeletonized neurons downloaded from the FlyWire Codex website (version 783 of the FAFB dataset). To obtain consistent scaling for the computation of fractal dimension, we chose to normalize the coordinates in the skeleton. We performed min-max scaling, to contain the skeleton in the unit cube while preserving its aspect ratio.

Splitting was performed using synapse flow centrality metric which determines the maximum flow/number of paths between pre- and postsynapses in the skeletonized neurons. The skeleton is split into compartments at the most distal nodes/points with maximal centrality ^29^.

Here we noticed that for some neurons, multiple dendritic and/or axonal arbors were found. However, in many of these cases multiple arbors were relatively short (one example shown in Supplementary Figure S1.1.). In order to avoid these small arbors affecting our statistics, we decided to work only with the largest arbor, defined by the number of nodes in the tree representation of the neuron.

### Geometric properties

The total cable length (i.e., the total physical length of the neuron) was directly available from the dataset and therefore did not require additional computation. In contrast, estimating the fractal dimension required a substantially more complex analysis.

### Estimating fractal dimension

Since the Drosophila connectome contains an enormously large number of neurons, our calculations needed to be as general as possible. For the estimation, we used a simple box-counting method ^30^. Using the numpy.logspace function, we generated box sizes between *log*_10_ (*w*) and *log*_10_ (1), where *w* is the maximum physical distance between two neighboring nodes in the skeleton.

We discovered in the dataset that there are few exceptional cases where some errors happened during the skeletonization of the reconstructed neurons, resulting in nodes missing and the neighbors being connected with unrealistic long lines (Supplementary Figure S1.2.). Consequently, the automated fractal dimension estimation for these neurons yields values that are excessively large or small and not physically meaningful. To enable reliable computation of fractal dimension, we decided to apply some thresholding: neurons were excluded if the maximum distance between any two neighboring nodes exceeded 25% of the total span (the distance between the two furthest points in the neuron skeleton). This threshold was chosen because it is approximately the distance where the distribution changes its shape indicating that these irregularities appear (Supplementary Figure S1.3).

After applying this filtering procedure, we analyzed a total of 125330 neurons. Even with the previously mentioned corrections, we were not able to filter out all cases in which the estimated fractal dimension was below 1. Previously it already has been discussed how non-linear box-counting plots can appear for neurons ^5^. We noticed some similar effects in our analysis, with constant parts right around the limits/boundaries of the curve (Supplementary Figure S1.4.). To address this, we excluded the plateau regions and restricted our linear regression strictly to the intermediate scaling range, bounded by the inner edges of the plateaus, to determine the corrected fractal dimension. However, for a subset of extremely sparse neurons, this correction was insufficient, and the fractal dimension remained below 1.0. To resolve this, we interpolated between the existing nodes of these sparse neurons to generate a higher-density spatial representation, and then recalculated their fractal dimension.

In the same iteration, we also implemented a minor correction at the neuron level. In cases where the estimated fractal dimension was below 1, we interpolated between all neighboring nodes, by inserting extra nodes with equal distances, thereby limiting the maximum distance to the value used for interpolation. We were aiming to have a relatively small distance between the neighboring nodes, without sacrificing runtime. Finally we chose a distance of 0.02% of the maximum distance of the total span. We were hoping to partially resolve the problem of the splitting providing extra small axonal and dendritic parts, because this is when these problems occur. However, there are still some axons and dendrites with fractal dimensions below 1. These usually have very small cable length and maximum width, meaning these are small, highly elongated structures where the branching density is low (Supplementary Figure S1.5).

### Preparing the data for topological properties

Once the data were loaded, neurons were prepared for graph-based analysis of tree properties. Neuronal skeletons often have multiple nodes between two branching nodes in order to follow the neuron’s curvature. To remove these, the neurons were downsampled using the navis.downsample_neuron function and ensuring that all intermediate nodes were removed, to reduce node density along branches while preserving branching structure and connectivity. This step minimizes the influence of sampling resolution and geometric path length, allowing subsequent analyses to focus on topological complexity rather than absolute neurite length.

### Topological properties

To characterize neuronal morphology, we analyzed neurons based on their topological as well as geometric properties. For the graph-based analysis, we computed the total number of nodes, tree height, and maximum tree width (defined as the maximum number of nodes at the same depth), see Supplementary Fig. S1.6. All measurements were performed on the prepared NetworkX graph representations of the neurons.

### Comparing distributions of neuron properties

Cliff’s δ is a non-parametric effect size metric used to measure the amount of differences between two groups of single-variable observations ^32^. It is calculated as the probability of values in one distribution being larger than in another one minus the reverse probability:

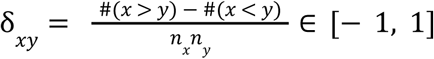

where *x* and *y* are the variables from the distributions; *n_x_* and *n_y_* are the sample sizes; # represents counting operations. The Cliff’s δ values are bounded in the [-1, 1] interval, with the sign indicating the direction of the shift. Different significance thresholds can be defined based on the magnitude ^33^: |δ| < 0. 11 is insignificant; 0. 11 < |δ| < 0. 28 is small; 0. 28 < |δ| < 0. 43 is medium; |δ| > 0. 43 indicates large effect size.

### Center of mass for each class of neurons

In order to visualize better the clusters on correlation maps, we computed the center of masses for each class of neurons using the formulas: 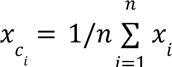 and 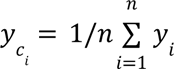. These are shown as larger colored circles on the correlation maps.

## Data & code availability

The connectome data were downloaded through the FlyWire project’s Codex web interface and fafbseg.py python API.

The analysis was performed by a set of commercially available python packages and custom code. Neuron morphological properties, such as the cable length of different compartments were obtained through *navis*. The neuron tree’s topological properties were computed using the *networkx* library. All the custom codes written for this analysis are available at https://github.com/BeaMarton13/DrosophilaFractalNeurons2026.

Fractal dimension was computed by a custom script that uses multiple standard python packages: *pandas*, *numpy*, *scipy*, *scikit-learn*. For visualization, the following packages were used: *matplotlib*, *fafbseg*, *caveclient*.

## Acknowledgements

This study was supported by the Romanian National Authority for Scientific Research and Innovation, CNCS-UEFISCDI, projects: PN-IV-P6-6.1-CoEx-2024-0139 (M.E-R), ERANET-NEURON-2-IBRAA (M.E-R), FLAGERA-JTC2023-MONAD (M.E-R), PN-IV-P2–2.1-TE-2023-1548 (M.E-R), PN-IV-P2-2.1-TE-2023-0558 (M.E-R); and by the Romanian Ministry of Research, Innovation and Digitalization, project HUNTGEN nr. PNRR-III-C9-2022-I8-66 (I-B. M.). B.P. was also supported by the Collegium Talentum Program of Hungary.

## Author contributions

Ildikó-Beáta Márton: Software; Data curation; Formal analysis; Investigation; Methodology; Validation; Visualization; Writing – original draft; Writing – review & editing. Balázs Péntek: Data curation; Formal analysis; Investigation; Methodology; Software; Validation; Visualization; Writing – original draft; Writing – review & editing. Mária Ercsey-Ravasz: Conceptualization; Funding acquisition; Investigation; Methodology; Project administration; Supervision; Writing – original draft; Writing – review & editing.

