## Supplementary Information for "The trade-off between fractal morphology and function of Drosophila neurons"

**1. Methods** 1
**2. Results** 5
**2.1. Intrinsic neurons** 5
**2.2. Afferent and efferent neurons** 8
**2.3. Afferent sensory neurons** 13
**References** 17

### 1. Methods

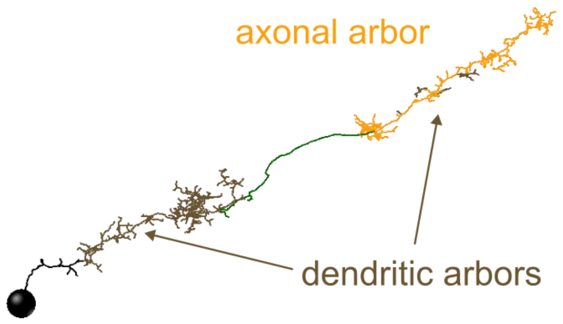


Supplementary Figure S1.1: **Example of a neuron skeleton after splitting.** Resulting in multiple dendritic abors.

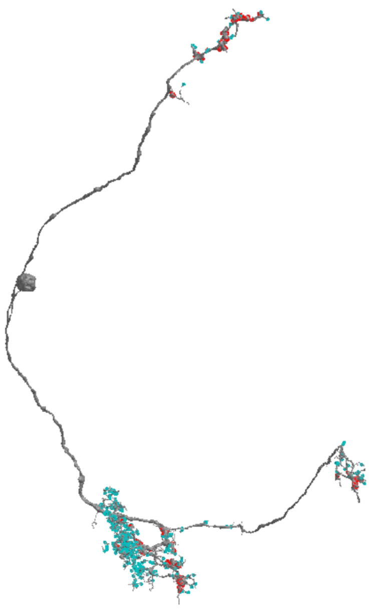

Supplementary Figure S1.2: **Example for a full neuron before skeletonization (left) and after skeletonization and automatic healing (right).**

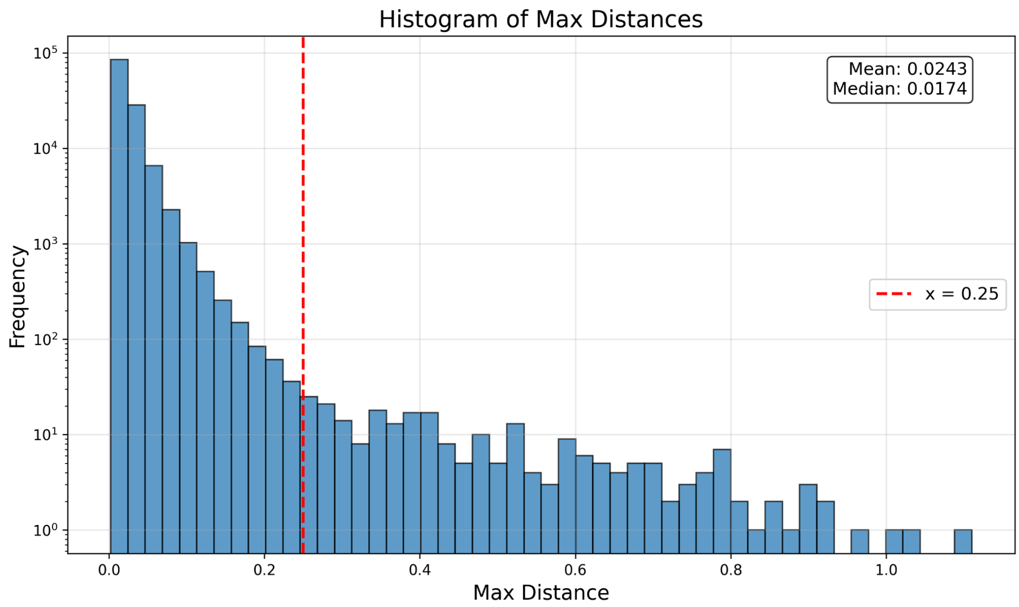


Supplementary Figure S1.3: **Histogram of maximum distances.** Maximum distances of neighboring nodes within each neuron skeleton.

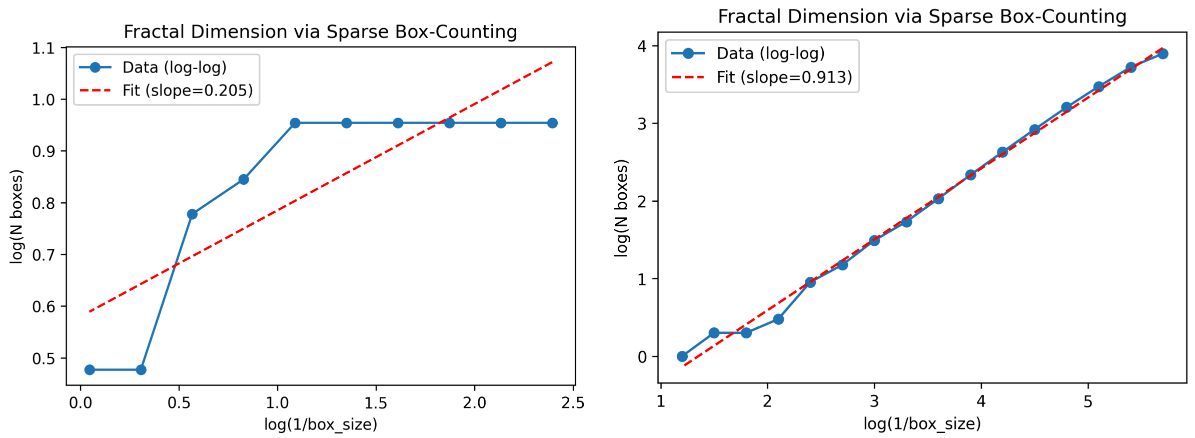


Supplementary Figure 1.4: **Log-log plot of box count versus inverse box size for box-counting analysis.** The blue points represent measured data, and the red dashed line shows the linear fit with slope ~0.205. Note that the first fit is performed over the full range of data points, including the constant (plateau) region at small inverse box sizes, which pulls the estimated slope downward. The second fit, restricted to the non-constant (scaling) region alone, yields a steeper slope and a more accurate estimate of the fractal dimension. In our analysis, the fitting is only done on this non-constant part.

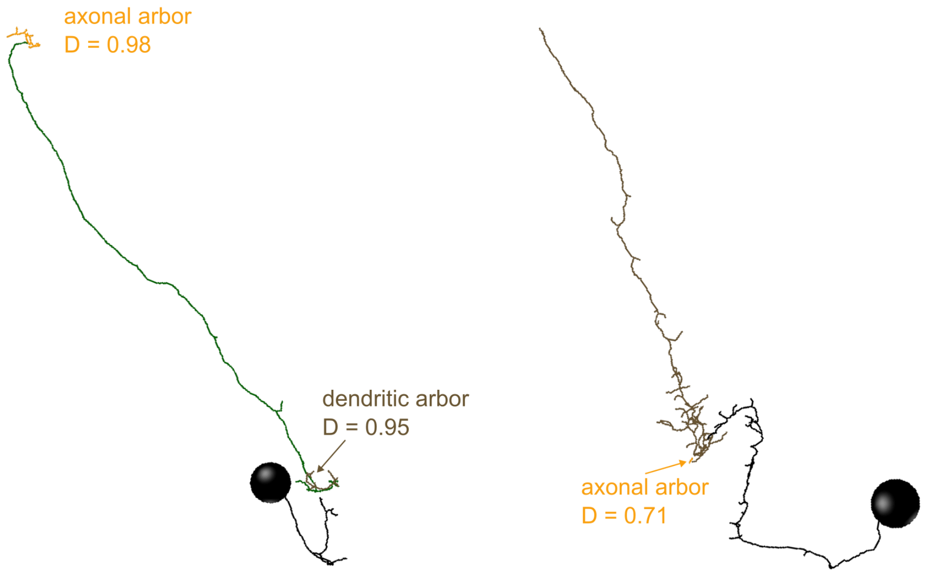


Supplementary Figure S1.5: **Example of two neuron skeletons with arbors providing fractal dimensions smaller than 1**. This happens when these arbors are extremely small and simple, normally the dimension should be 1, just the automatized algorithm and the fitting procedures fail because of the small number of available points.

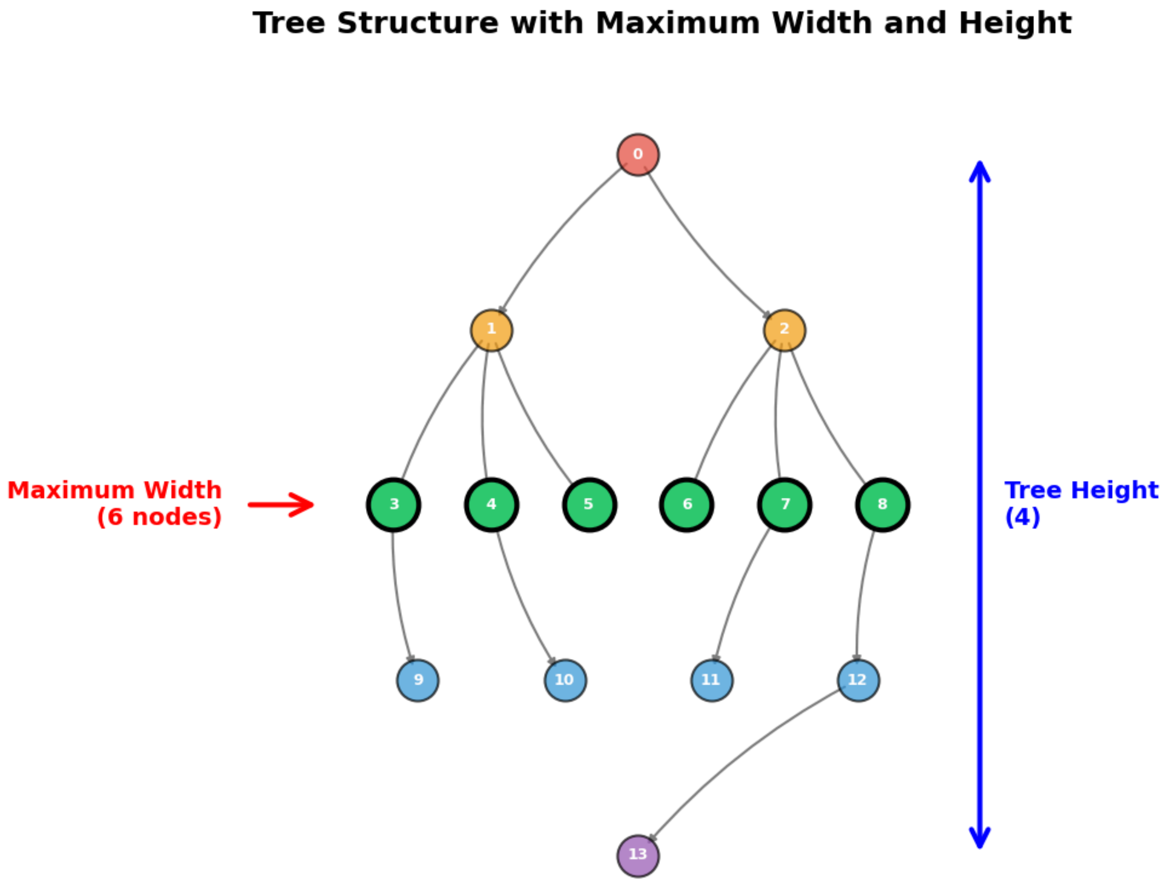


Supplementary Figure S1.6: **Example of a tree structure illustrating key topological properties.**

### 2. Results

#### 2.1. Intrinsic neurons

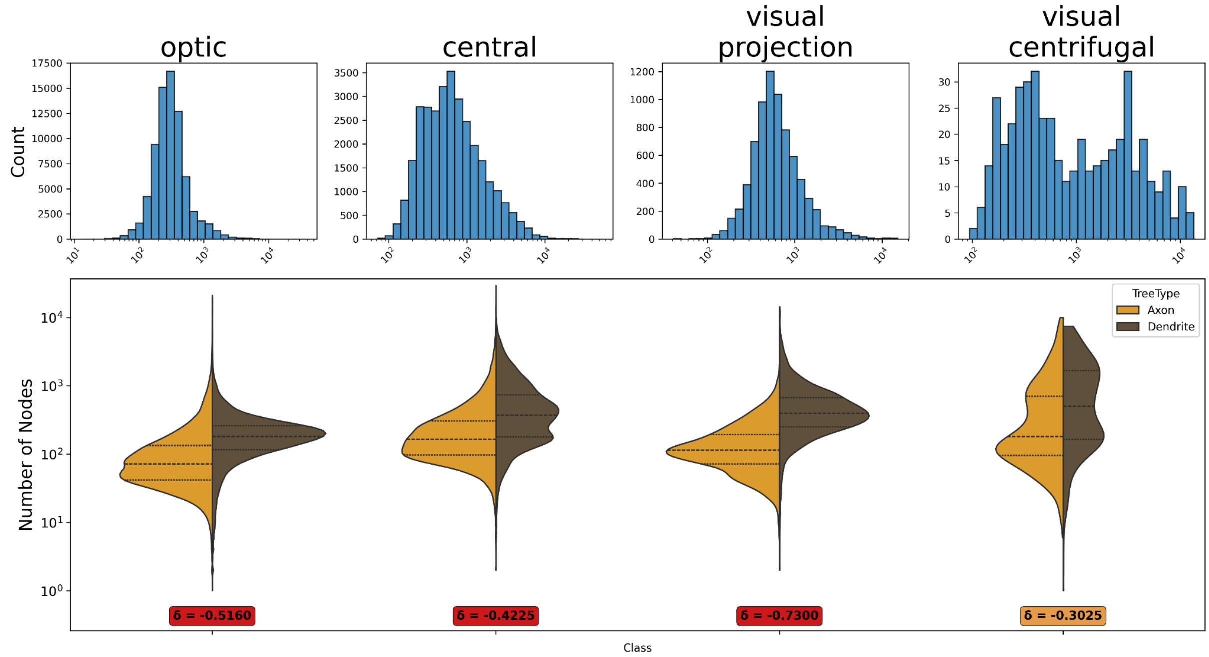


Supplementary Figure S2.1: **Violin plots of number of nodes in neuron trees, shown for intrinsic super-classes.** Top row: histograms for full neurons; bottom row: distributions for axonal/dendritic arbors.

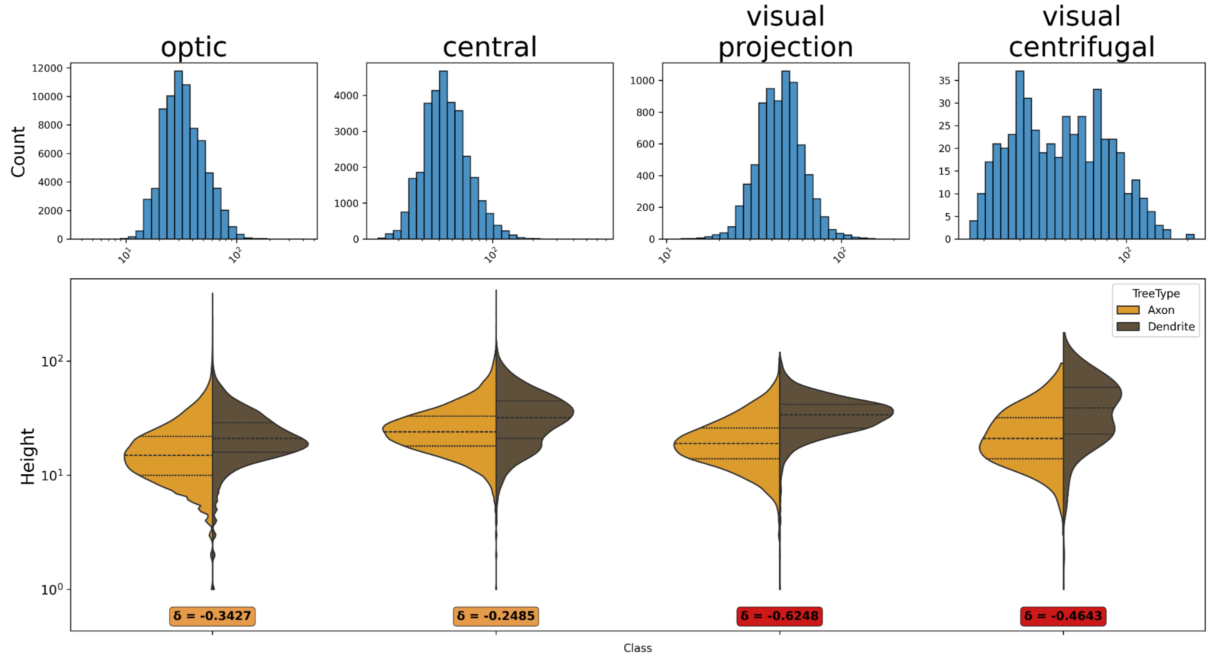


Supplementary Figure S2.2: **Violin plots of tree height, shown for intrinsic super-classes.** Top row: histograms for full neurons; bottom row: distributions for axonal/dendritic arbors.

| Comparison of dendritic vs. axonal arbor properties: intrinsic neuron super-classes | | |
|---|---|---|
| Super Class | Property | Cliff's δ |
| Optic | Cable Length | -0.4129 |
| | Fractal Dimension | -0.4819 |
| | Number of Nodes | -0.5160 |
| | Maximum Width | -0.5150 |
| | Tree Height | -0.3427 |
| Central | Cable Length | -0.2730 |
| | Fractal Dimension | -0.3042 |
| | Number of Nodes | -0.4225 |
| | Maximum Width | -0.4523 |
| | Tree Height | -0.2485 |
| Visual Projection | Cable Length | -0.5399 |
| | Fractal Dimension | -0.6587 |
| | Number of Nodes | -0.7300 |
| | Maximum Width | -0.6997 |
| | Tree Height | -0.6248 |
| Visual Centrifugal | Cable Length | -0.0773 |
| | Fractal Dimension | -0.0339 |
| | Number of Nodes | -0.3025 |
| | Maximum Width | -0.1599 |
| | Tree Height | -0.4643 |

Supplementary Table S2.1: **Comparison of dendritic vs. axonal arbor properties: intrinsic neuron super-classes.** Colored based on Cliff's δ significance thresholds[1]: insignificant (<0.11) - white, small (>0.11) - green, medium (>0.28) - orange and large (>0.43) - red.

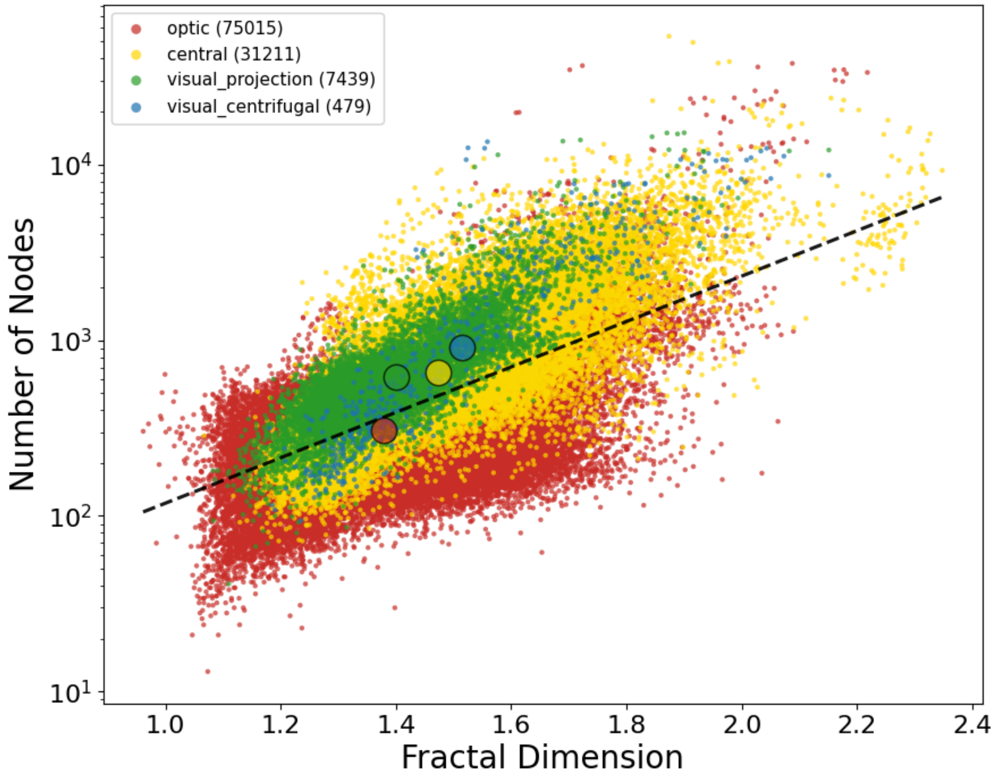


Supplementary Figure S2.3: **Colored correlation plot.** Fractal dimension vs number of nodes for intrinsic super-classes.

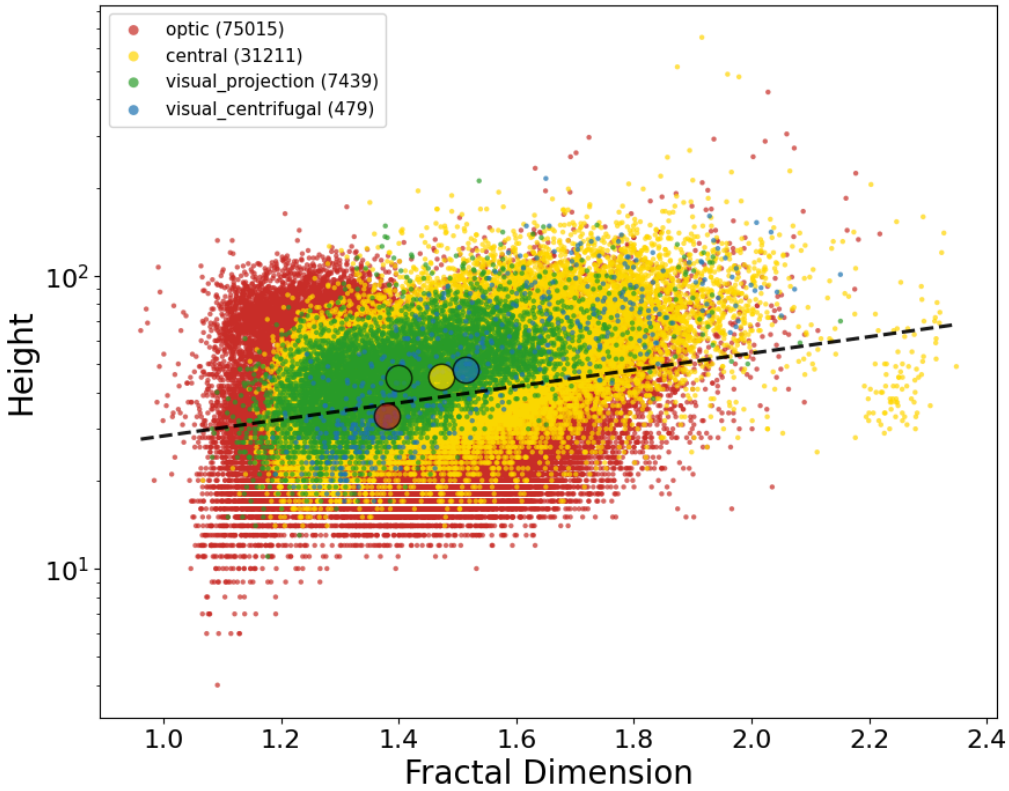


Supplementary Figure S2.4: **Colored correlation plot.** Fractal dimension vs height for intrinsic super-classes.

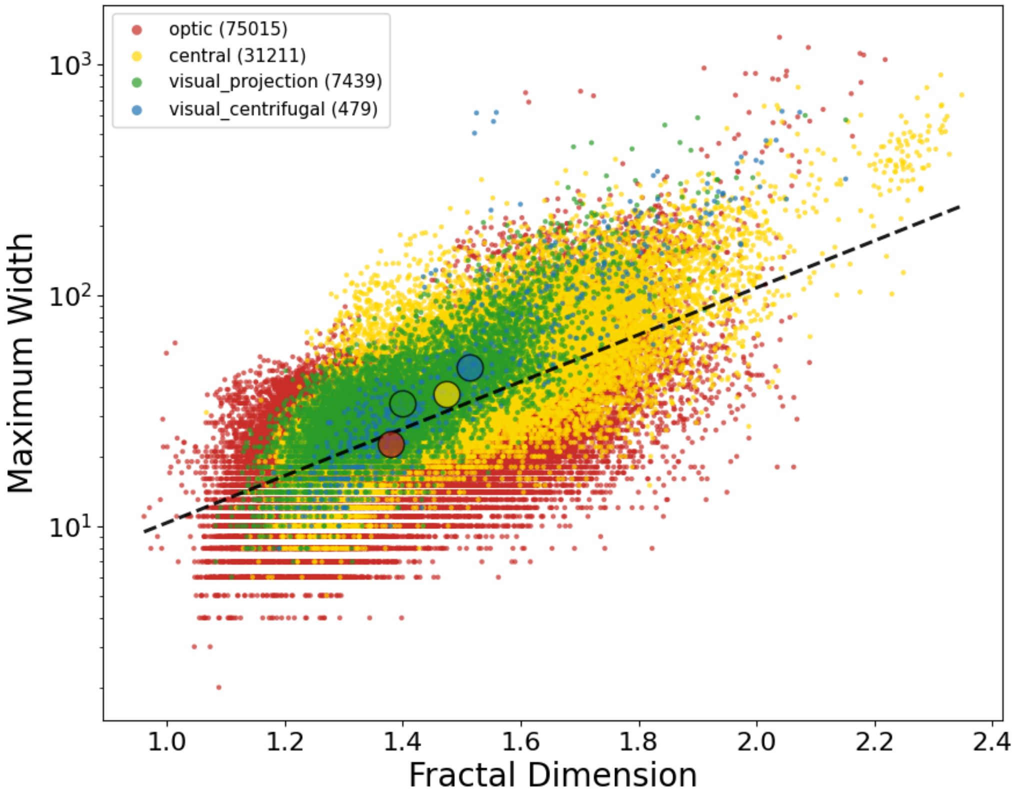


Supplementary Figure S2.5: **Colored correlation plot.** Fractal dimension vs maximum width for intrinsic super-classes.

#### 2.2. Afferent and efferent neurons

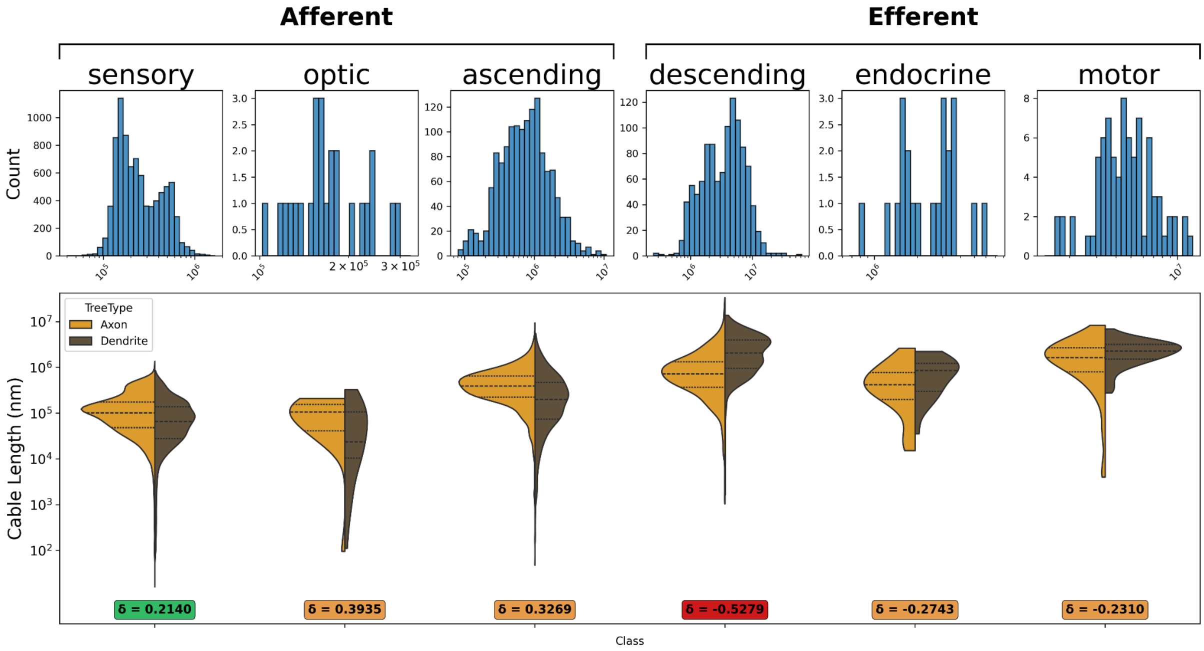


Supplementary Figure S2.6: **Violin plots of cable length for afferent and efferent super-classes.** Top row: histograms for full neurons; bottom row: distributions for axonal/dendritic arbors.

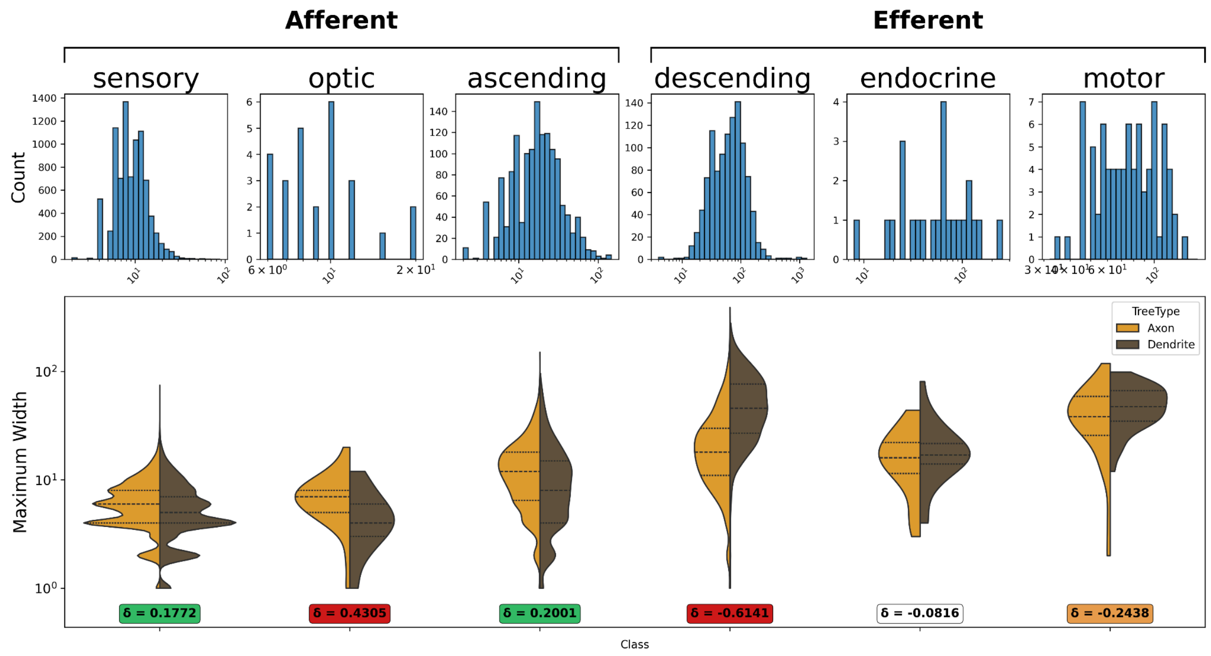


Supplementary Figure S2.7: **Violin plots of maximum width for afferent and efferent super-classes.** Top row: histograms for full neurons; bottom row: distributions for axonal/dendritic arbors.

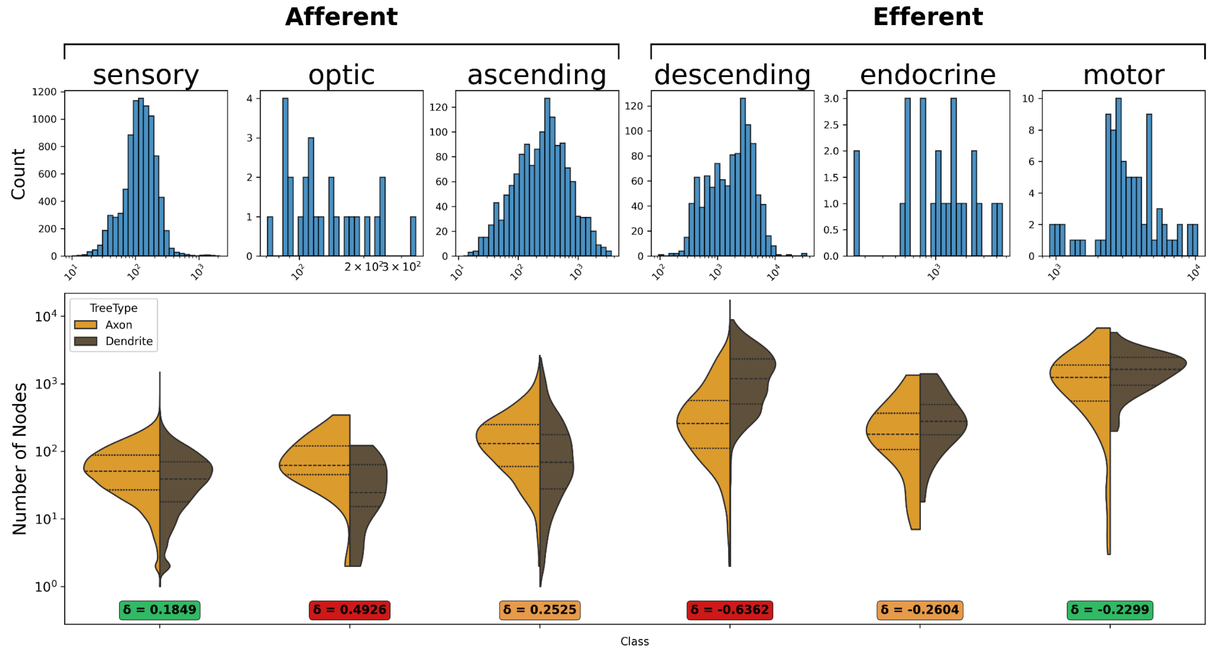


Supplementary Figure S2.8: **Violin plots of number of nodes for afferent and efferent super-classes.** Top row: histograms for full neurons; bottom row: distributions for axonal/dendritic arbors.

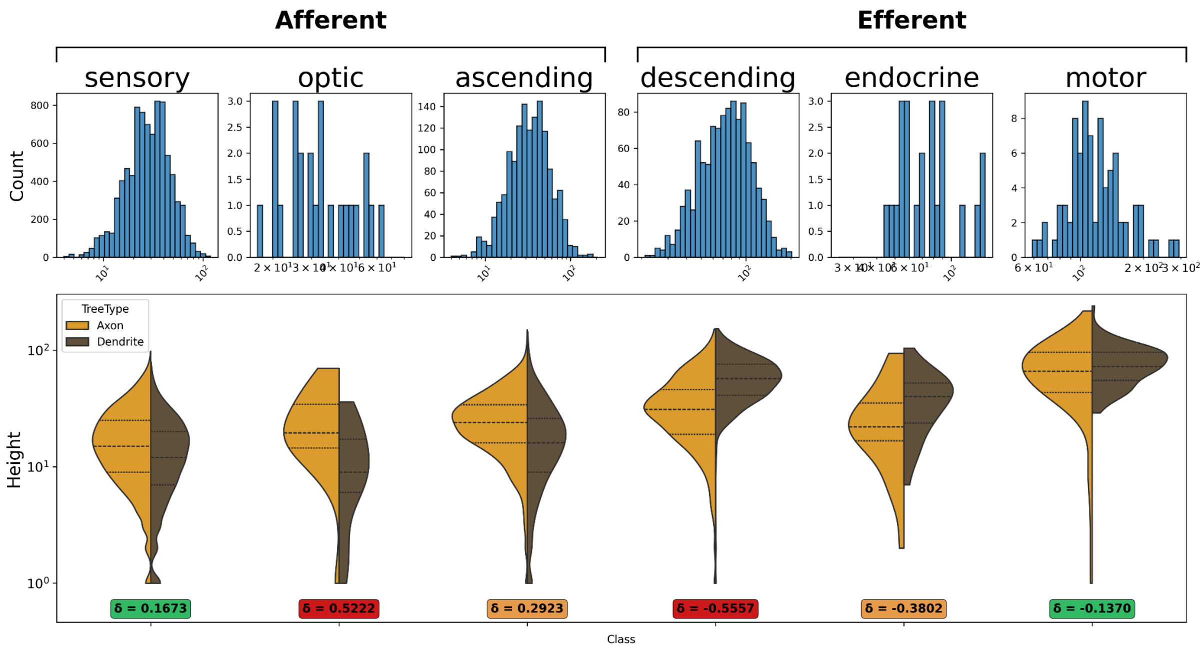


Supplementary Figure S2.9: **Violin plots of height for afferent and efferent super-classes.** Top row: histograms for full neurons; bottom row: distributions for axonal/dendritic arbors.

| Comparison of dendritic vs. axonal arbor properties: afferent/efferent neuron super-classes | | |
|---|---|---|
| **Super Class** | **Property** | **Cliff's δ** |
| Sensory | Cable Length | 0.2140 |
| | Fractal Dimension | 0.1544 |
| | Number of Nodes | 0.1849 |
| | Maximum Width | 0.1772 |
| | Tree Height | 0.1673 |
| Ascending | Cable Length | 0.3269 |
| | Fractal Dimension | 0.2298 |
| | Number of Nodes | 0.2525 |
| | Maximum Width | 0.2001 |
| | Tree Height | 0.2923 |
| Descending | Cable Length | -0.5279 |
| | Fractal Dimension | -0.6327 |
| | Number of Nodes | -0.6362 |
| | Maximum Width | -0.6141 |

| Comparison of dendritic vs. axonal arbor properties: afferent/efferent neuron super-classes | | |
|---|---|---|
| | Tree Height | -0.5557 |
| Endocrine | Cable Length | -0.2743 |
| | Fractal Dimension | -0.0694 |
| | Number of Nodes | -0.2604 |
| | Maximum Width | -0.0816 |
| | Tree Height | -0.3802 |
| Optic | Cable Length | 0.3935 |
| | Fractal Dimension | 0.4541 |
| | Number of Nodes | 0.4926 |
| | Maximum Width | 0.4305 |
| | Tree Height | 0.5222 |
| Motor | Cable Length | -0.2310 |
| | Fractal Dimension | -0.2132 |
| | Number of Nodes | -0.2299 |
| | Maximum Width | -0.2438 |
| | Tree Height | -0.1370 |

Supplementary Table S2.2: **Comparison of dendritic vs. axonal arbor properties: afferent/efferent neuron super-classes.** Colored based on Cliff's δ significance thresholds[1]: insignificant (<0.11) - white, small (>0.11) - green, medium (>0.28) - orange and large (>0.43) - red.

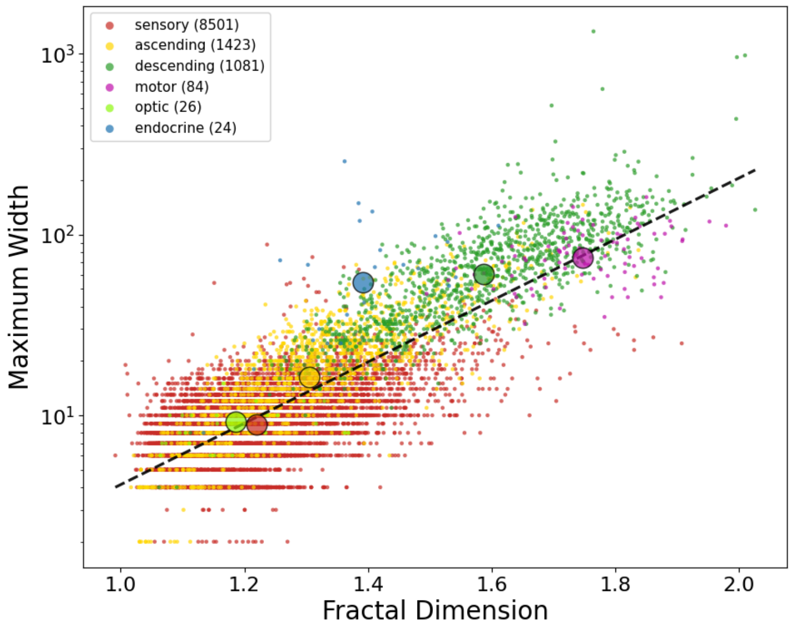


Supplementary Figure S2.10: **Colored correlation plot.** Fractal dimension vs maximum width for afferent and efferent super-classes.

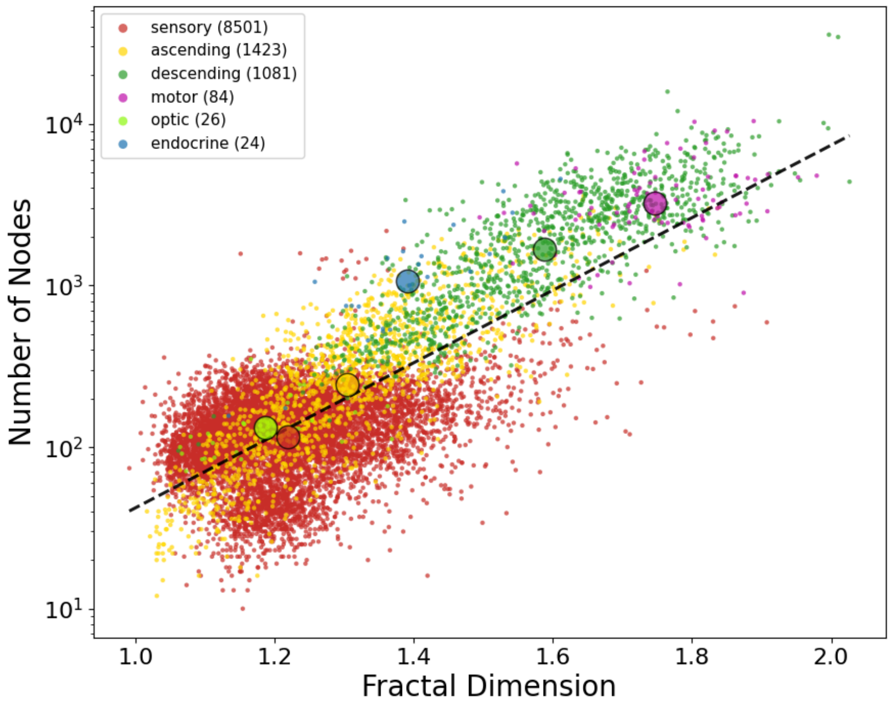


Supplementary Figure S2.11: **Colored correlation plot.** Fractal dimension vs number of nodes for afferent and efferent super-classes.

#### 2.3. Afferent sensory neurons

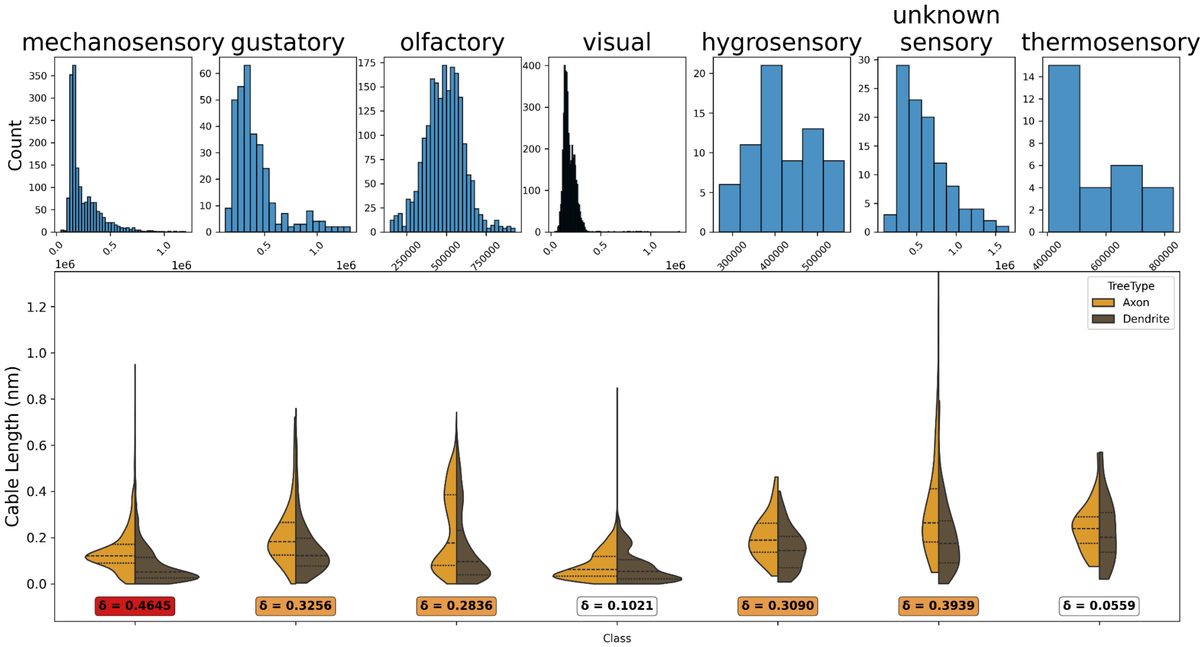


Supplementary Figure S2.12: **Violin plots of cable length for sensory classes.** Top row: histograms for full neurons; bottom row: distributions for axonal/dendritic arbors.

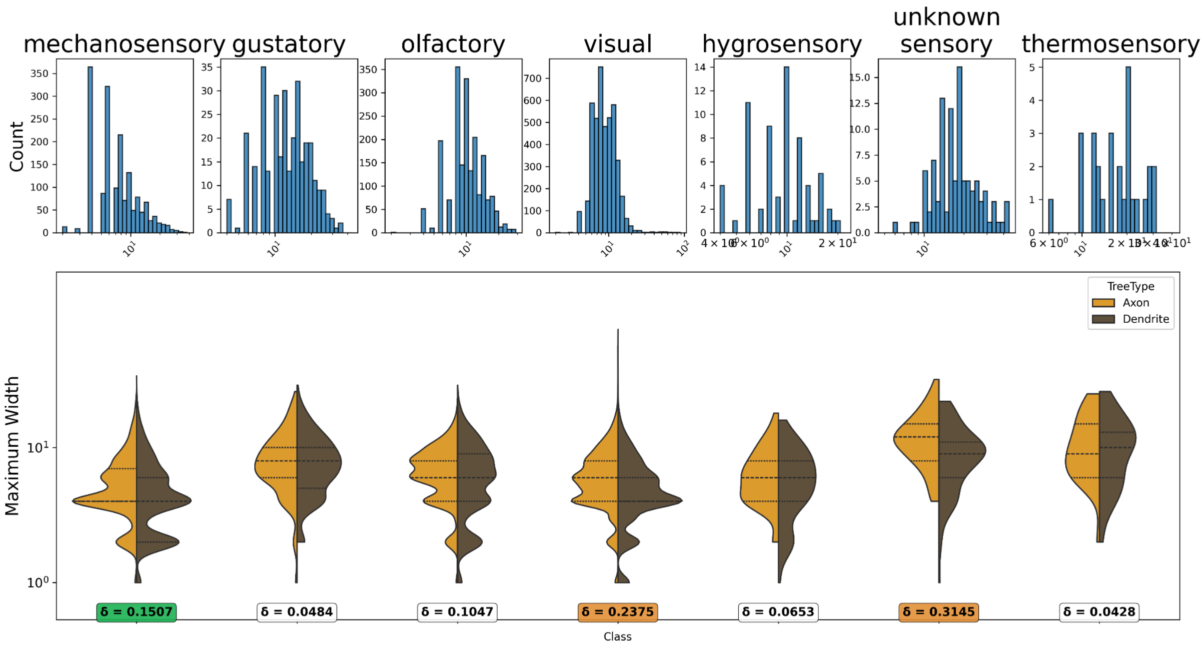


Supplementary Figure S2.13: **Violin plots of maximum width for sensory classes.** Top row: histograms for full neurons; bottom row: distributions for axonal/dendritic arbors.

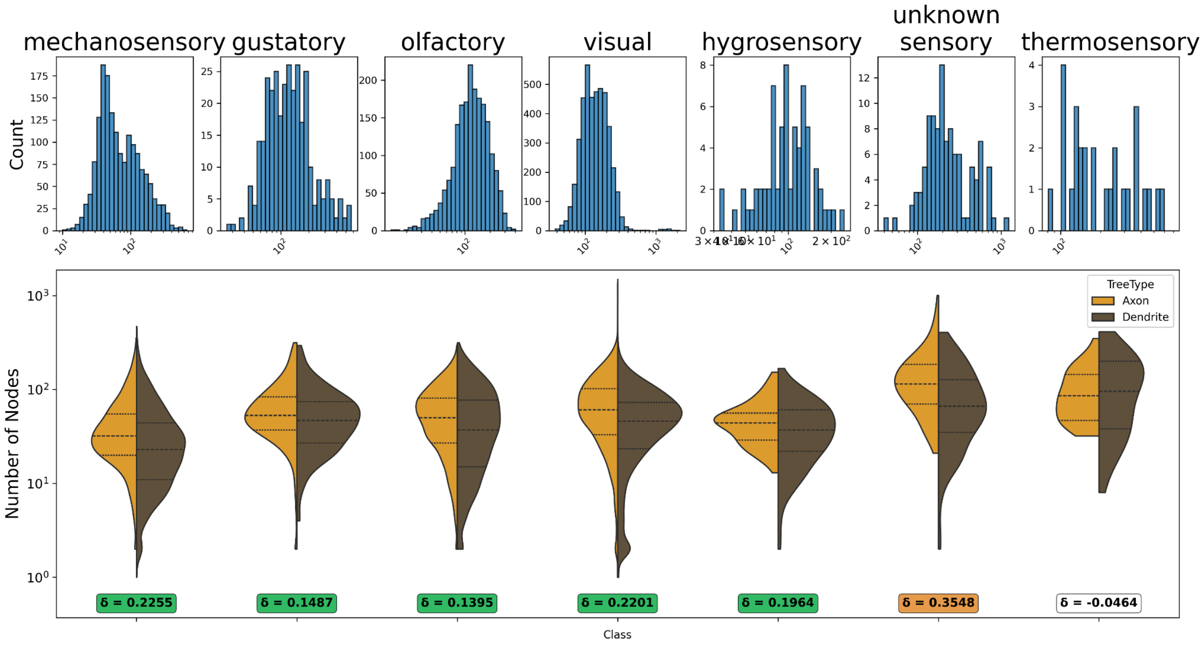


Supplementary Figure S2.14: **Violin plots of number of nodes for sensory classes.** Top row: histograms for full neurons; bottom row: distributions for axonal/dendritic arbors.

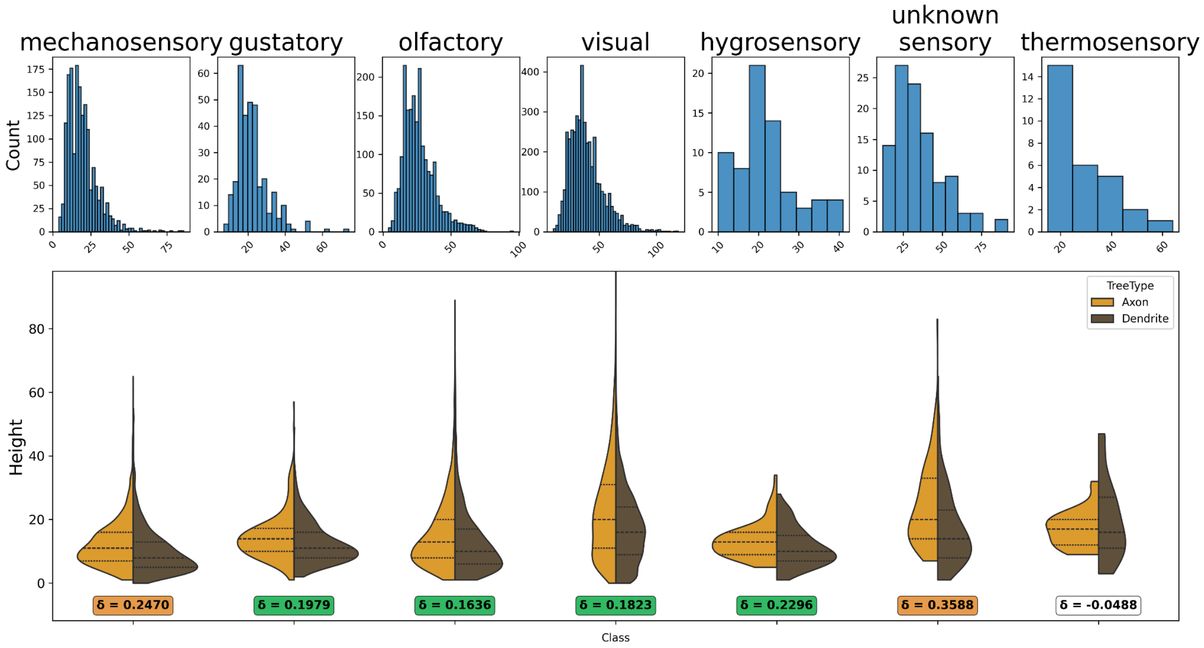


Supplementary Figure S2.15: **Violin plots of height for sensory classes.** Top row: histograms for full neurons; bottom row: distributions for axonal/dendritic arbors.

| Comparison of dendritic vs. axonal arbor properties: sensory neuron classes | | |
|---|---|---|
| Class | Property | Cliff's δ |
| Mechanosensory | Cable Length | 0.4645 |
| | Fractal Dimension | 0.1459 |
| | Number of Nodes | 0.2255 |
| | Maximum Width | 0.1507 |
| | Tree Height | 0.2470 |
| Gustatory | Cable Length | 0.3256 |
| | Fractal Dimension | 0.0629 |
| | Number of Nodes | 0.1487 |
| | Maximum Width | 0.0484 |
| | Tree Height | 0.1979 |
| Olfactory | Cable Length | 0.2836 |
| | Fractal Dimension | 0.1933 |
| | Number of Nodes | 0.1395 |
| | Maximum Width | 0.1047 |
| | Tree Height | 0.1636 |
| Visual | Cable Length | 0.1021 |
| | Fractal Dimension | 0.1420 |
| | Number of Nodes | 0.2201 |
| | Maximum Width | 0.2375 |
| | Tree Height | 0.1823 |
| Hygrosensory | Cable Length | 0.3090 |
| | Fractal Dimension | 0.1464 |
| | Number of Nodes | 0.1964 |
| | Maximum Width | 0.0653 |
| | Tree Height | 0.2296 |
| Unknown Sensory | Cable Length | 0.3939 |
| | Fractal Dimension | 0.3350 |

| Comparison of dendritic vs. axonal arbor properties: sensory neuron classes | | |
|---|---|---|
| | Number of Nodes | 0.3548 |
| | Maximum Width | 0.3145 |
| | Tree Height | 0.3588 |
| Thermosensory | Cable Length | 0.0559 |
| | Fractal Dimension | -0.0083 |
| | Number of Nodes | -0.0464 |
| | Maximum Width | 0.0428 |
| | Tree Height | -0.0488 |

Supplementary Table S2.3: **Comparison of dendritic vs. axonal arbor properties: sensory neuron classes.** Colored based on Cliff's δ significance thresholds[1]: insignificant (<0.11) - white, small (>0.11) - green, medium (>0.28) - orange and large (>0.43) - red.

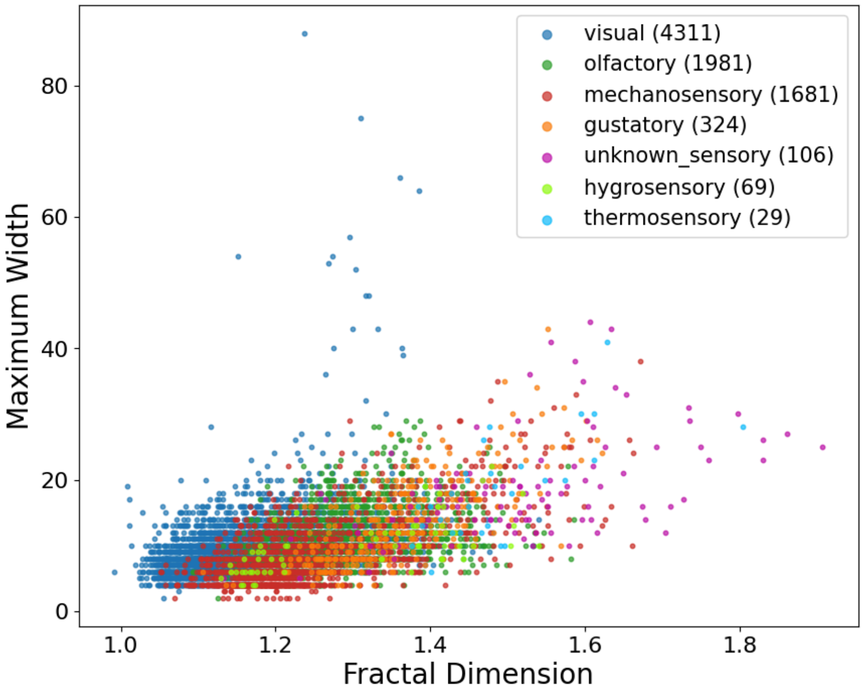


Supplementary Figure S2.16: **Colored correlation plots.** Fractal dimension vs maximum width for sensory classes.

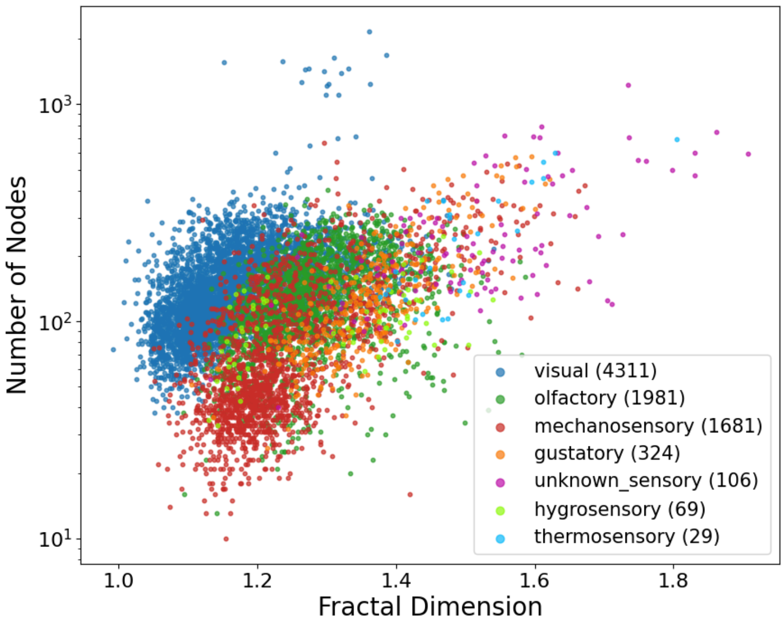


Supplementary Figure S2.17: **Colored correlation plots.** Fractal dimension vs number of nodes for sensory classes.
